# Structure-Based Virtual Screening Identifies TREM2-Targeted Small Molecules that Enhance Microglial Phagocytosis

**DOI:** 10.1101/2025.08.23.671917

**Authors:** Sungwoo Cho, Baljit Kaur, Kevin Lam, Farida El Gaamouch, Katarzyna Kuncewicz, Niklas Piet Doering, Gerhard Wolber, Moustafa T. Gabr

## Abstract

Triggering receptor expressed on myeloid cells 2 (TREM2) is a microglia-specific receptor whose activation promotes phagocytosis and neuroprotection in Alzheimer’s disease (AD) and related neurodegenerative disorders. While therapeutic efforts have largely focused on antibodies, small molecule TREM2 modulators remain limited. Here, we applied a structure-based virtual screening workflow targeting a putative allosteric site on TREM2, guided by PyRod-derived pharmacophores from molecular dynamics simulations. Screening of the Enamine Collection yielded 20 candidate compounds, three of which demonstrated binding in TRIC assays. The top hit, **EN020**, exhibited a KD of 14.2 µM (MST) and 35.9 µM (SPR), and significantly enhanced microglial phagocytosis in BV2 cells outperforming the known TREM2 agonist **VG-3927**. A preliminary structure–activity relationship (SAR) study, including synthetic and catalog-derived analogs, highlighted a narrow tolerance for scaffold modifications, with only **T2V002** retaining partial TREM2 binding affinity. This work identifies **EN020** as a novel small molecule TREM2 modulator with functional activity, providing a framework for rational optimization toward potential AD therapeutics.

## Introduction

Alzheimer’s disease (AD) remains a major unmet medical challenge, with available therapies offering only symptomatic relief and failing to stop or meaningfully delay disease progression.^[^¹,²^]^ Hallmark pathological features of AD include extracellular deposits of amyloid-β (Aβ) peptides, intracellular neurofibrillary tangles composed of hyperphosphorylated tau, and a sustained neuroinflammatory state.^[^³⁻⁵^]^ Neuroinflammation, defined as inflammatory activity within the central nervous system (CNS),^[^⁵^]^ is largely orchestrated by microglia, the brain’s innate immune cells, which, upon activation, secrete a variety of proinflammatory mediators.^[^⁶^]^ Persistent overproduction of these factors fosters a neurotoxic milieu, exacerbating neuronal damage and accelerating disease advancement.^[^⁵^]^ Traditional therapeutic development has primarily targeted cholinergic neurotransmission, β- secretase activity, Aβ aggregation, or tau pathology. In contrast, recent efforts increasingly highlight the modulation of microglial inflammatory responses as a compelling and innovative strategy for AD treatment.^[7,8]^

Triggering receptor expressed on myeloid cells 2 (TREM2) is a membrane receptor uniquely enriched in microglia within the central nervous system (CNS).^[9]^ Upon ligand engagement, TREM2 signals through the adaptor protein DAP12 — encoded by TYROBP and containing an immunoreceptor tyrosine-based activation motif (ITAM) — to regulate essential microglial activities such as phagocytosis, survival, proliferation, motility, and lysosomal function.^[9–12]^ The discovery that rare TREM2 variants are strong genetic risk factors for AD has spurred extensive investigation into its contribution to disease pathogenesis.^[13,14]^ In both amyloidogenic mouse models and AD patients carrying the R47H variant, loss of TREM2 function compromises microglial recruitment and clustering around Aβ plaques, resulting in less compact, more diffuse deposits. This deficiency also undermines microglial survival during reactive microgliosis, heightens susceptibility to Aβ-mediated toxicity, and aggravates neuronal injury.^[13,14]^ Transcriptomic analyses reveal that TREM2-deficient microglia fail to acquire a protective gene expression program; although changes in overall Aβ load vary depending on model and disease stage, increased neuritic degeneration is a consistent finding.^[13,15]^ Conversely, overexpression of human TREM2 enhances microglial phagocytic capacity, upregulates related gene networks, and reduces both amyloid burden and neuritic pathology.^[16]^ Similarly, agonistic antibodies targeting TREM2 have been shown to foster plaque compaction, lower Aβ levels, attenuate neuritic injury, and improve cognitive and behavioral outcomes in mouse models.^[17,18]^

Most therapeutic efforts to modulate TREM2 activity have centered on antibody-based agents, whereas small-molecule strategies remain comparatively underexplored. Antibodies for Alzheimer’s disease, while effective in target engagement, face inherent challenges such as limited penetration across the blood–brain barrier (BBB), high production costs, risk of eliciting immune responses, and prolonged systemic half-lives that can extend immune-related adverse events.^[19,20]^ Small molecules, by contrast, present several attractive features: they typically achieve superior BBB permeability, can be administered orally, carry a lower likelihood of immunogenicity, and allow for precise pharmacokinetic (PK) optimization to enable adjustable dosing and mitigate side effects.^[21,22]^ Moreover, their comparatively low manufacturing costs could improve patient accessibility.^[22]^ Incorporating small molecules into the AD immunotherapy landscape may therefore accelerate progress, particularly in combination regimens that target multiple immune receptors. In this vein, our group recently identified **MG-257** as the first small molecule reported to influence the interaction between TREM2 and galectin-3.^[23]^ More recently, **VG-3927** emerged as the first reported small molecule TREM2 agonist with therapeutic potential for AD.^[24]^ This compound exhibits high potency, robust brain penetration, and the ability to induce anti-inflammatory microglial activation, attenuate AD-associated pathology in humanized mouse models, and demonstrate confirmed CNS activity in non-human primates.^[24]^ In this study, we applied a structure-based virtual screening approach to identify small-molecule modulators of TREM2, guided by PyRod^[25]-^derived pharmacophores from molecular dynamics simulations. Hits from the Enamine Screening Collection were prioritized for biophysical evaluation, followed by structure–activity relationship (SAR) exploration and functional assessment of their ability to enhance microglial phagocytosis.

## Results and Discussion

### Generation of the 3D Pharmacophore for Virtual Screening

To develop a 3D pharmacophore for virtual screening, PyRod^[25]^ was used to identify potential hotspots for small molecule intermolecular interactions with the protein. PyRod traces water molecules in several short molecular dynamics simulations and derives dynamic molecular interaction fields (dMIFs) for chemical feature recognition. The Ig-like domain of human TREM2 (PDB: 5UD7)^[26]^ was searched for suitable binding pockets for targeting. One potential allosteric binding site was identified on the opposite side of the complementarity-determining region (CDR), responsible for binding endogenous ligands. The cavity is formed by β-sheet C, C’, C’’, D, E, the connecting loop between E and F; and β-sheet G **(Figure 1).**

**Figure. 1.**
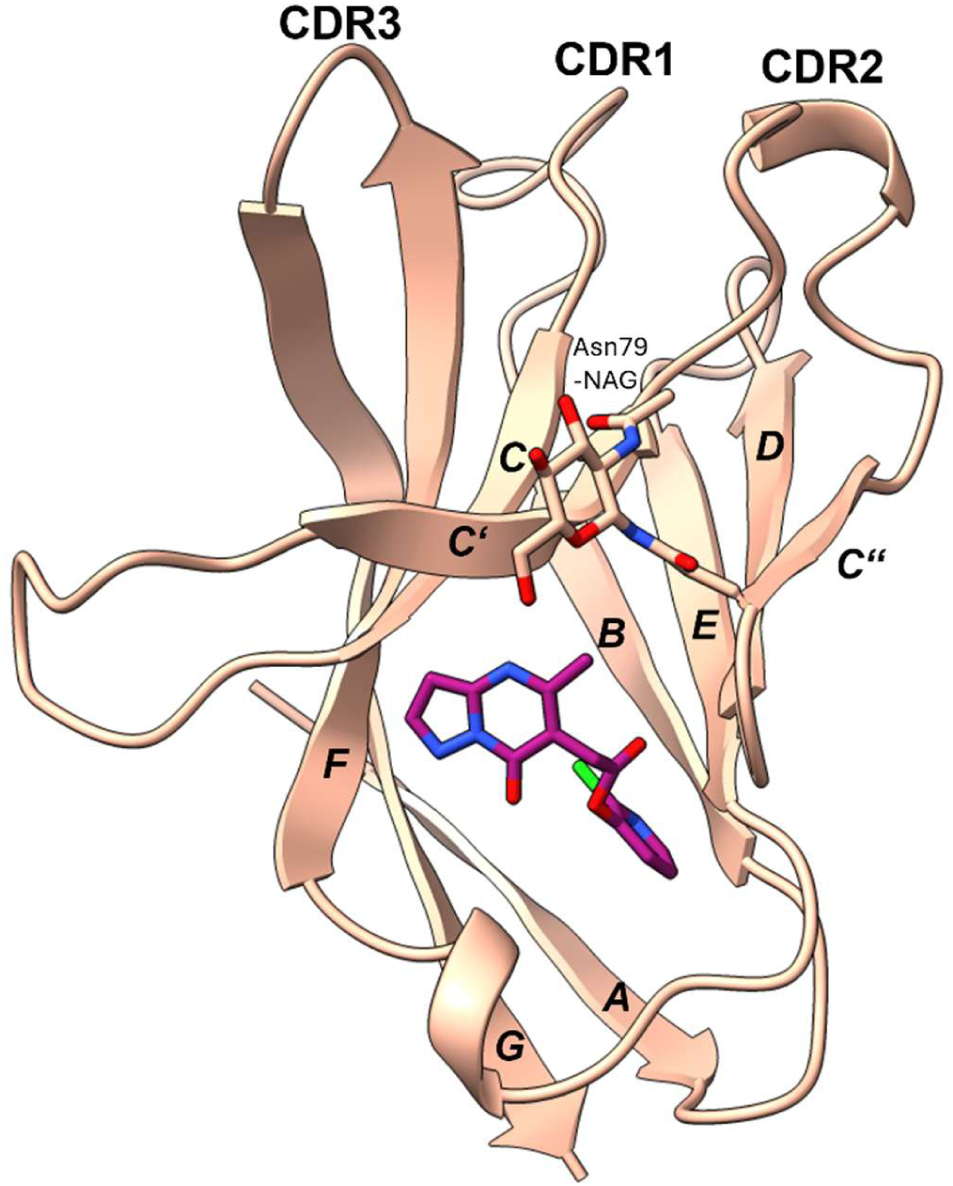
Crystal structure of the Ig-like domain of human TREM2 (PDB: 5UD7) with EN020 docked into the allosteric binding site.

To generate the dMIFs in the selected cavity, MD simulations were conducted and submitted to PyRod^[25]^ **(Figure 2)**. Generated dMIFs revealed three hydrophobic hotspots formed by (i) Trp50, Leu97, Tyr108 deep in the cavity, (ii) between Val63, Val64, Thr82, and Asn79-NAG, and (iii) between Gln61 and Val63. Three hotspots for hydrogen acceptors were identified between Trp50, Leu97, Tyr108: one close to Arg52, and one between Ser81 and Thr82. One large hotspot for hydrogen donor bonds surrounds Asp104, and two smaller ones close to Tyr108 and Thr82.

**Figure 2.**
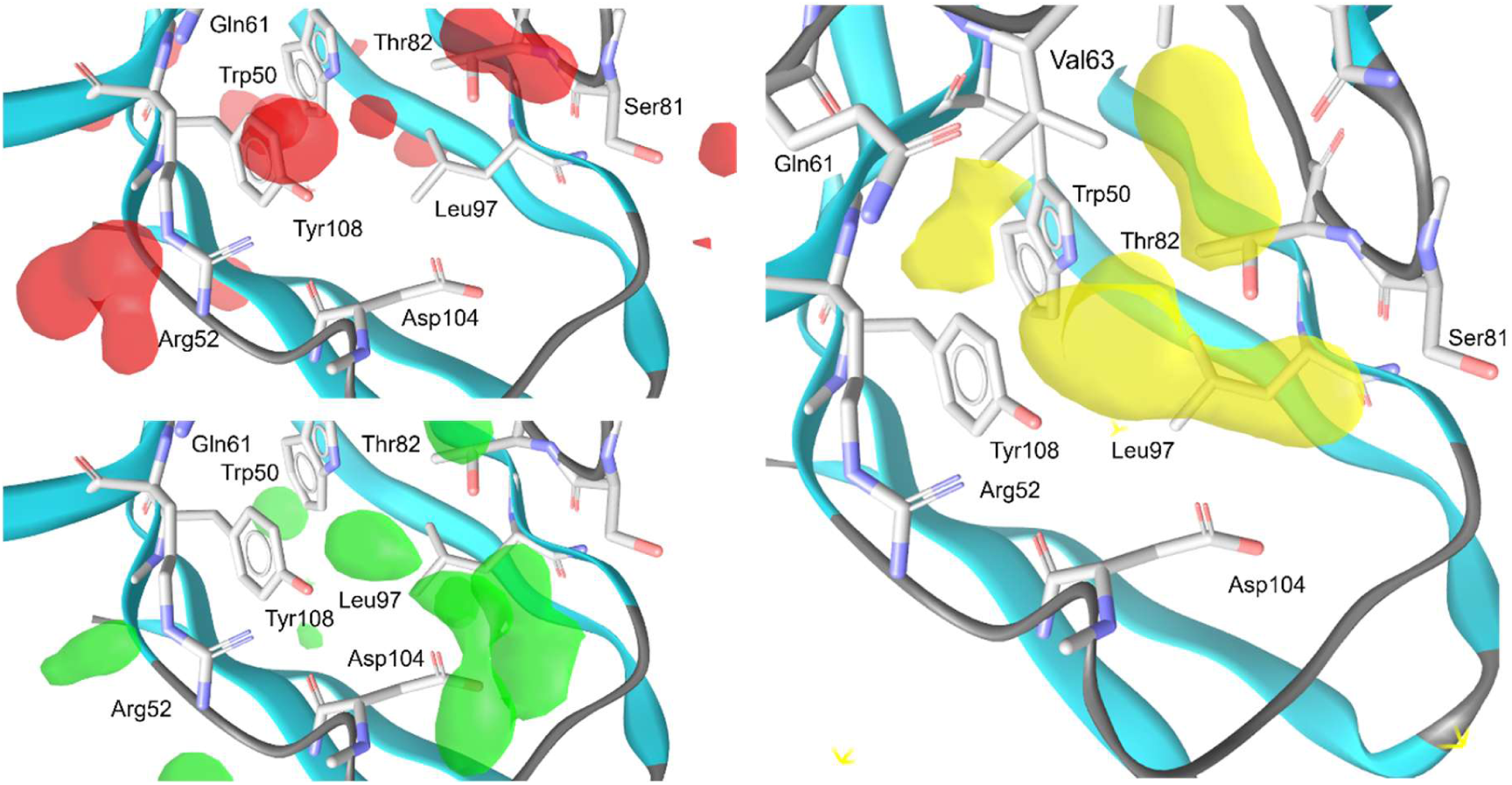
**Dynamic interaction fields (dMIFs) of the proposed TREM2 binding cavity generated by PyRod**^[25]^, indicating hotspots for specific interactions. Red fields for hydrogen bond acceptors, green for hydrogen bond donors and yellow for hydrophobic contacts.

Pharmacophore features were then placed in these hotspots, favoring hydrophobic contacts favored due to their contribution to binding affinity.^[34]^ These steps resulted in one core pharmacophore (**Figure 3**) with four hydrophobic contacts, two of which are located deep in the cavity shaped by Trp50, Leu97, Leu100, Asp104, and Tyr108; two contacts on both sides of Val63; and a hydrogen acceptor bond with Arg52. It was also observed that at least one hydrogen acceptor bond near Ser81 or hydrogen donor bond near Asp104 or His103 improved the quality of the binding poses. Therefore, compounds had to fulfill at least one of the optional pharmacophore features, resulting in three sets of 3D pharmacophores.

**Figure 3.**
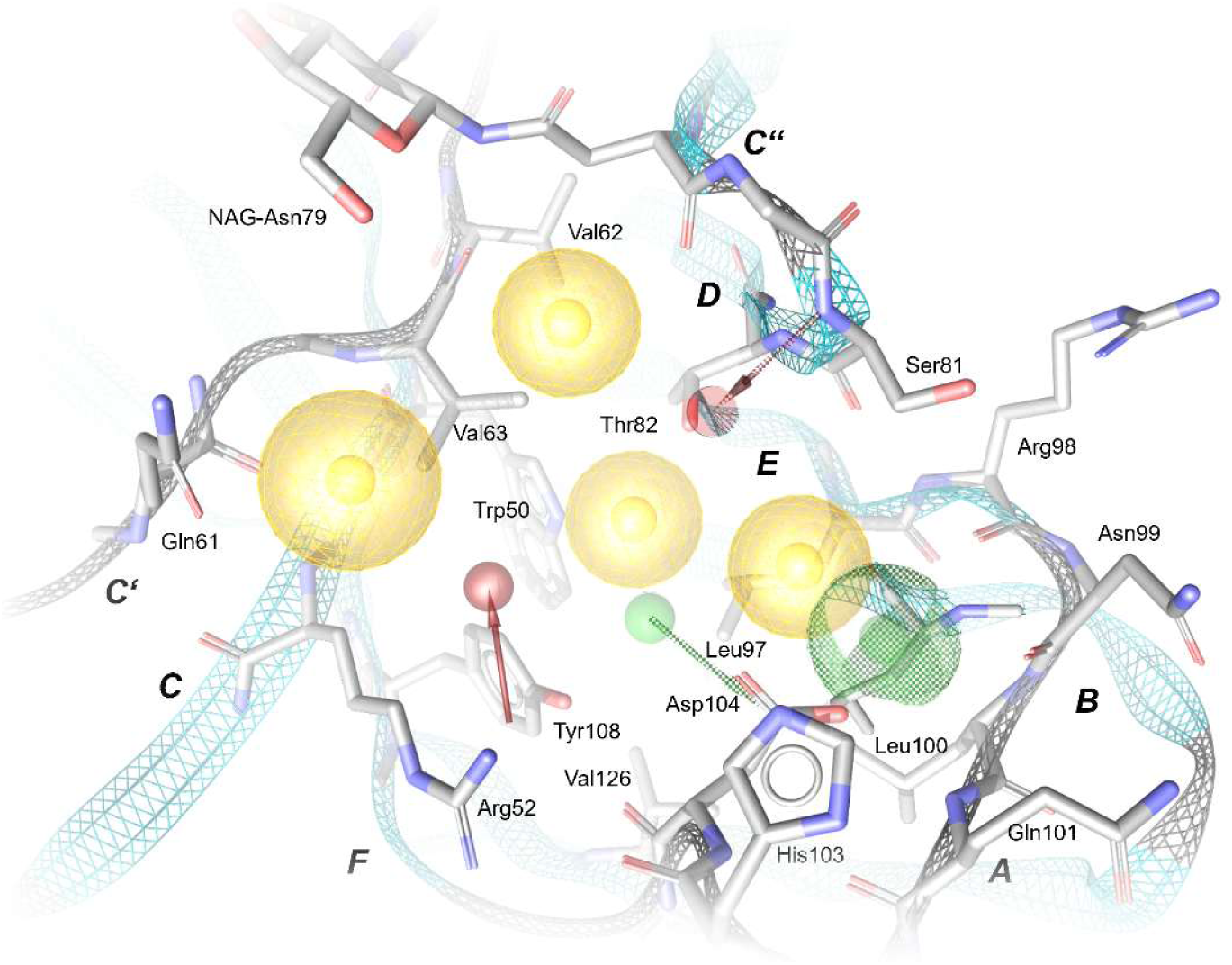
3D Pharmacophore for virtual screening of TREM2 with yellow spheres depicting hydrophobic contacts, red arrows as hydrogen bond acceptors and green arrow and spheres as hydrogen bond donors. Checkered pharmacophore features are optional. Compounds had to fulfill at least one of the optional pharmacophore features.

### Virtual Screening Campaign

The three sets of 3D pharmacophores were screened against the ESC and yielded 1312 hits overall. To validate the binding poses of the acquired hits, molecular docking was performed with each protonation state of the 1312 hits. For each protonation state ten binding hypotheses were built in the binding site and filtered for at least three hydrophobic contacts, which were then visually inspected. Binding hypotheses targeting the hydrophobic cavity close to Trp50, exhibiting favorable shape complementarity with the binding pocket and being able to match the core pharmacophore, were favored over the other. The visual inspection of the docking poses yielded twenty compounds (**Table S1**) for further experimental validation.

All binding poses of the selected compounds bind near the cavity shaped by Trp50, Leu97, Leu100, Asp104, and Tyr108 and fulfill the mandatory hydrogen acceptor bond with Arg50. They also form additional intermolecular interactions with the binding site to further increase binding affinity with TREM2. Energy minimization of the selected binding poses using the MMFF94 force field shows only minimal movement, indicating that the binding poses in TREM2 are close to their unbound conformation state.

### Allosteric Path Analysis

To further validate the selected binding pocket, we analyzed allosteric pathways to identify residues that may influence conformational changes across the transmembrane domain of TREM2 and thereby modulate its intracellular signaling. The analysis revealed two principal paths of allosteric communication flanking the predicted binding site. These paths converge at the base of the Ig-like domain, forming a single path that traverses the helical transmembrane domain and extends into the intracellular tail of TREM2 (**Figure 4A**). In particular, interactions involving Val62, Val63, and Trp50 appear to shape signaling from one side of the pocket, while Ser81, Leu100, and Val126 contribute from the opposite side (**Figure 4B**). These pathways were analyzed starting from the apo TREM2 structure and highlight the potential relevance of the identified binding pocket.

**Figure 4.**
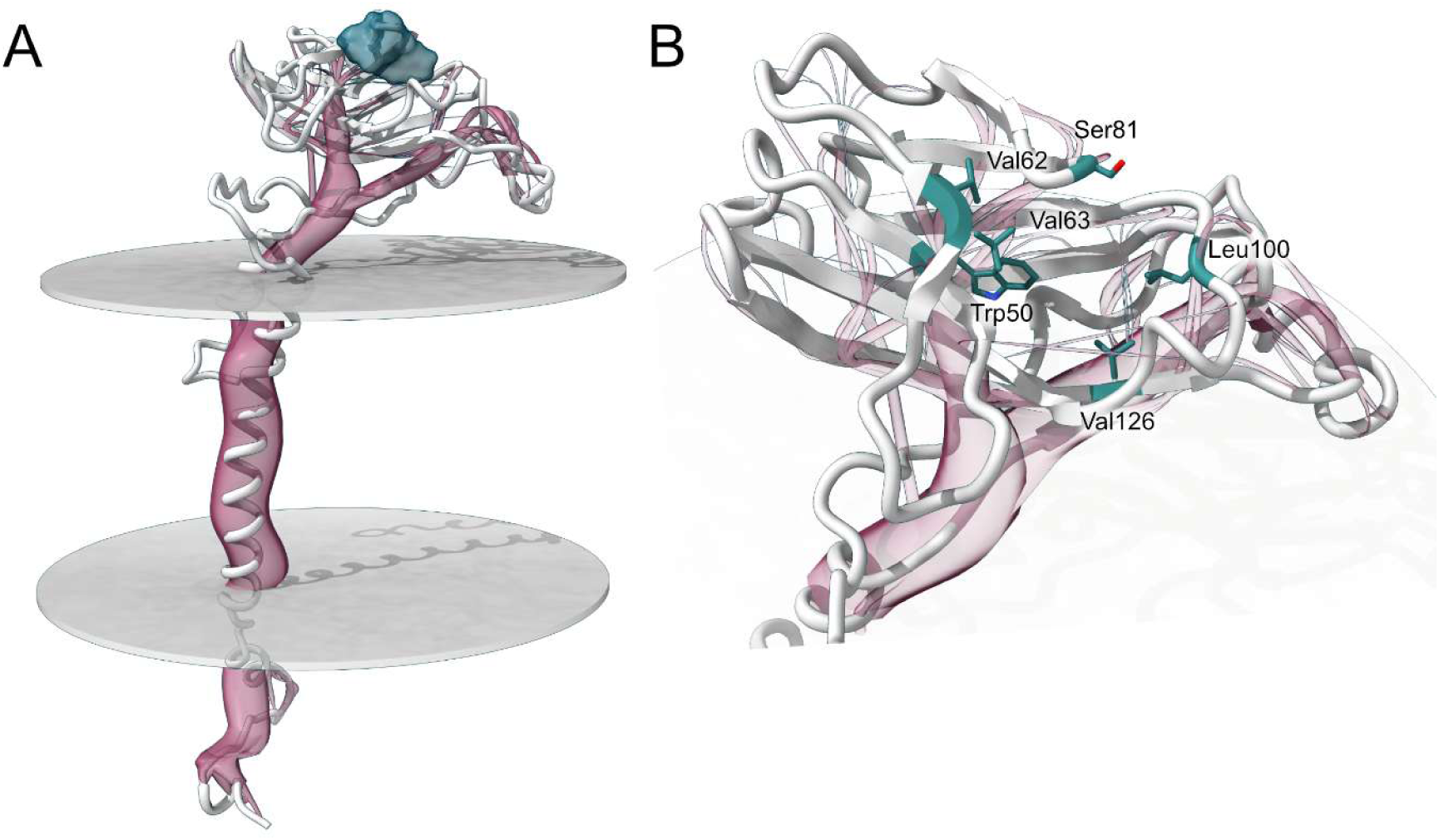
Allosteric path analysis of TREM2 conducted with MDPath. (A) Full allosteric paths (red) of TREM2. The paths originate around the predicted binding site (teal) and then traverse the transmembrane domain to the intracellular portion of TREM2. (B) A detailed view of the allosteric paths flanking the binding site. The residues from which the paths originate are highlighted in teal.

### Biophysical validation of TREM2 binding affinity

We performed single-dose screening using the Dianthus platform to evaluate 20 compounds derived from virtual screening hits with TREM2 protein at 100 μM using Temperature-Related Intensity Change (TRIC) analysis. Compounds showing signal changes greater than 5-fold the standard deviation of the negative control were defined as TREM2- binding hits. From this screening, we identified 3 out of 20 compounds as potential TREM2- binding hits (**Figure 5A**). To validate these hits, we conducted dose-response experiments for **EN007**, **EN017**, and **EN020** using TRIC-based binding assays (**Figure 5B-D**). Among the three candidates, only EN020 demonstrated clear concentration-dependent binding to TREM2 protein, while **EN007** and **EN017** showed no dose-dependent responses. Based on these results, we selected **EN020** as our primary TREM2-binding hit compound.

**Figure 5.**
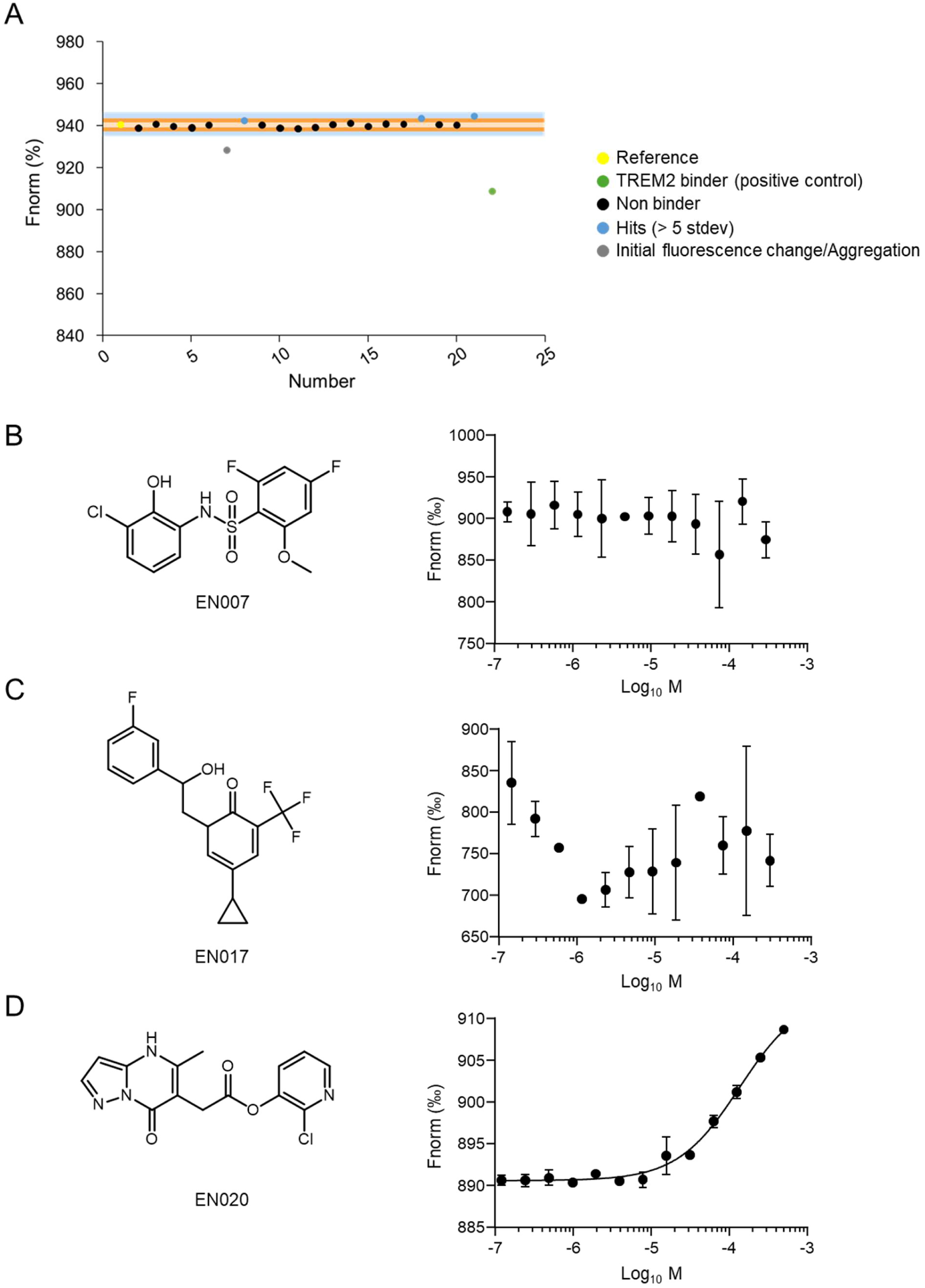
Identification and validation of TREM2-binding hits derived from Virtual Screening Hits**. A)** Single-concentration screening of 20 Virtual Screening Hits with TREM2 protein (10 nM) at 100 μM using TRIC analysis. Yellow dot: negative control (TREM2 alone); green dot: positive control (TREM2 with PC-192). Orange region: 3× standard deviation threshold; light blue region: 5× standard deviation threshold. Blue dots represent TREM2-binding hits above the threshold. Gray dots indicate false positives due to aggregation or fluorescence interference. **B-D)** Dose-response validation of three hit compounds (**EN007**, **EN017**, and **EN020**) using TRIC-based binding assays. Left panels show chemical structures of each compound; right panels show corresponding dose-response curves. Data are presented as mean ± SEM (n = 3).

### Biophysical validation of EN-20-TREM2 binding using MST and SPR

Based on the screening results from Dianthus, we utilized microscale thermophoresis (MST) and surface plasmon resonance (SPR) to further quantify the binding affinity of the identified hit compound **EN020** with TREM2. MST analysis confirmed the direct binding of **EN020** to TREM2 protein and revealed an KD of 14.24 ± 2.22 μM (**Figure 6A**). To obtain comprehensive binding kinetics data, we employed surface plasmon resonance (SPR) analysis, which enables real-time monitoring of molecular interactions and determination of both association (ka) and dissociation (kd) rate constants. In our experimental setup, recombinant TREM2 protein was immobilized onto the sensor chip surface, followed by injection of serial dilutions of **EN020** to generate concentration-dependent binding profiles. Real-time binding kinetics were captured as sensorgrams, demonstrating the interaction between **EN020** and immobilized TREM2 protein (**Figure 6B**). Through kinetic analysis of the binding curves, we determined that **EN020** exhibits a KD of 35.9 ± 7.13 μM for TREM2, confirming its direct interaction with the target protein and providing quantitative binding parameters for further compound optimization.

**Figure 6.**
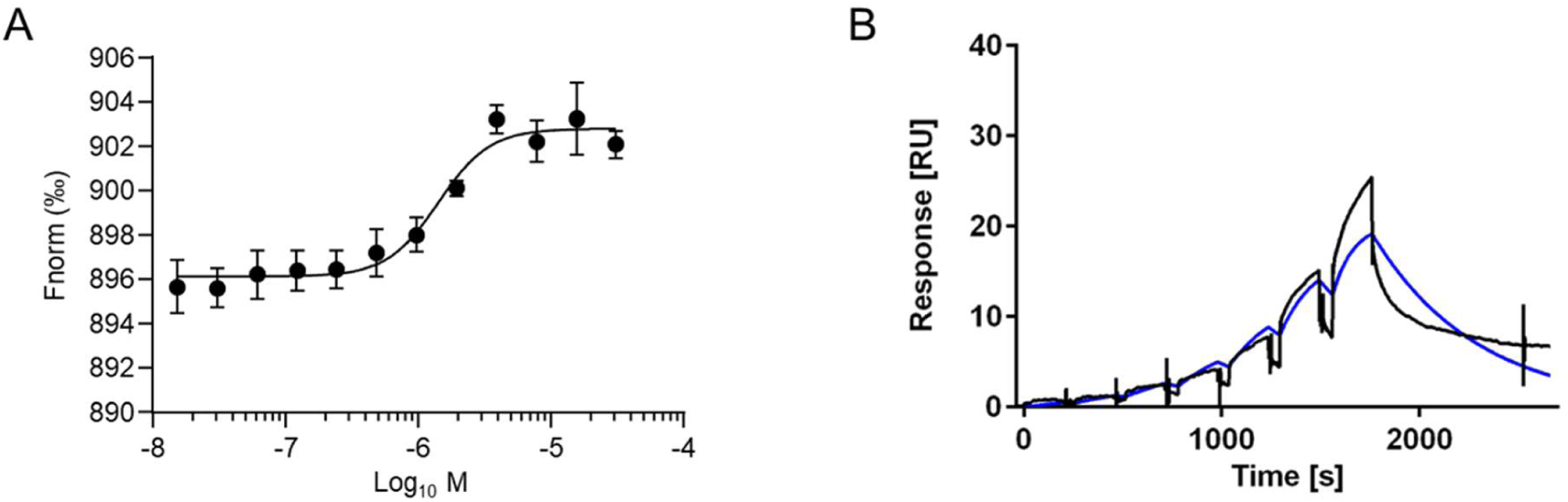
Identification and validation of TREM2 binding using MST and SPR. **A)** Dose-dependent binding experiments for **EN020** using MST. Data are presented as Mean ± SEM (n = 3). **B)** Single-cycle kinetic sensorgrams of **EN020** interacting with the TREM2 protein using SPR. Black lines represent the experimental data, while blue lines show the 1:1 kinetic binding model fit.

### Functional validation of EN020 as a TREM2 modulator via microglial phagocytosis assay

Microglial phagocytosis plays a pivotal role in maintaining brain homeostasis and limiting neurodegenerative processes by removing protein aggregates, apoptotic cells, and other debris. Dysregulation of this process is closely associated with AD and other neuroinflammatory disorders. To determine whether the TREM2-binding hit **EN020** could enhance microglial phagocytic function, we performed a bead uptake assay in BV2 murine microglial cells and compared its activity to that of **VG-3927**, a well-characterized small-molecule TREM2 agonist, and untreated controls. Cells were incubated with fluorescent latex beads following compound treatment, and the proportion of bead-positive cells was quantified. Both **VG-3927** and **EN020** increased the percentage of phagocytic cells relative to control, with **VG-3927** producing an approximate 36.5% increase and **EN020** eliciting nearly a 67% increase. The superior activity of **EN020** over **VG-3927** highlights its potential not only as a validated TREM2 binder but also as a functional enhancer of microglial activity, supporting its relevance for therapeutic development in neurodegenerative disease contexts **(Figure 7).**

**Figure 7.**
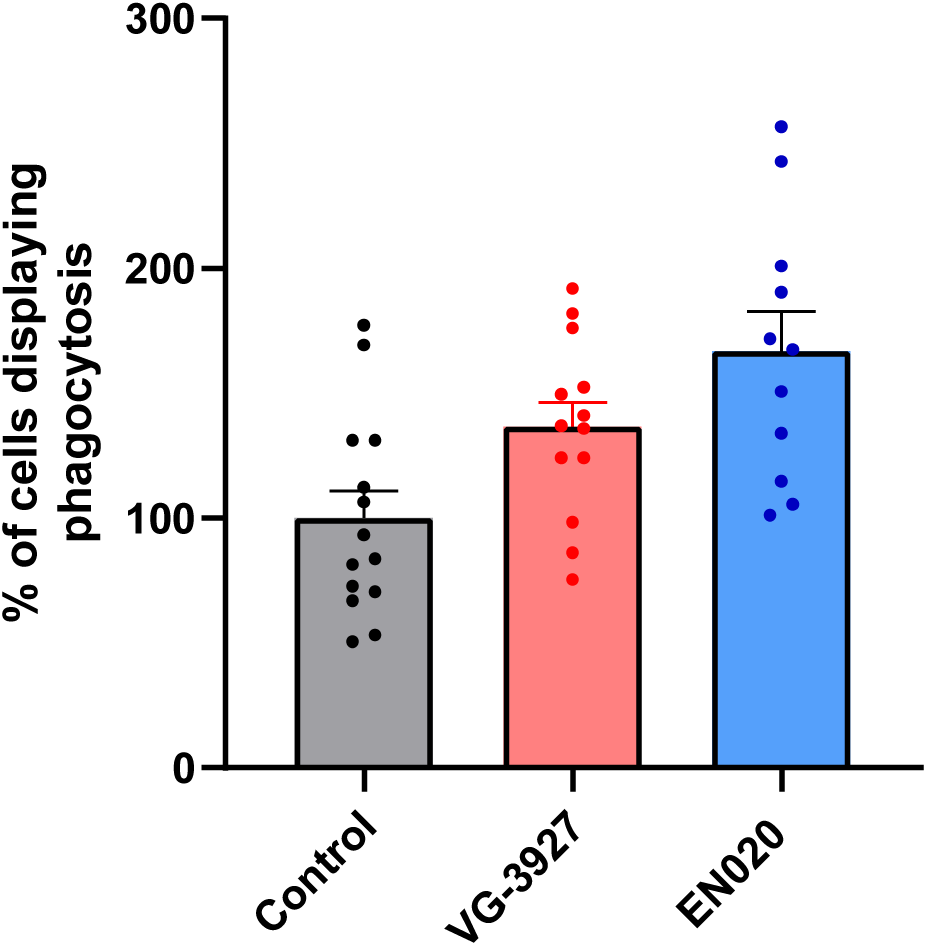
In vitro validation of the TREM2 binding activity. Histogram representing the quantification of BV2 cells showing at least one fluorescent latex bead in their cell body (percentage of control) after treatment with compounds **VG-3927** and **EN020** (25 μM-1 h) (n=5-8 pictures/group (N=2)).

### Binding mode of EN020 in TREM2

**EN020**’s binding mode was analyzed after molecular docking. **EN020** binds to the hydrophobic cavity with its chlorpyridin ring and fulfills all mandatory pharmacophore features of the PyRod^[25]^ pharmacophore **(Figure 8). EN020** forms four hydrophobic contacts with the binding site: chlorine with Val63, Thr82, Leu97 and Tyr108; the pyridine ring with Leu97; and the pyridine ring and the methyl moiety with Val63. The pyridine nitrogen forms the crucial hydrogen bond with Arg52. Additionally, the carbonyl oxygen of the ester moiety forms three hydrogen bond acceptors with the OH-moiety of Thr82 and the backbone of Ser81 and Thr82.

**Figure 8.**
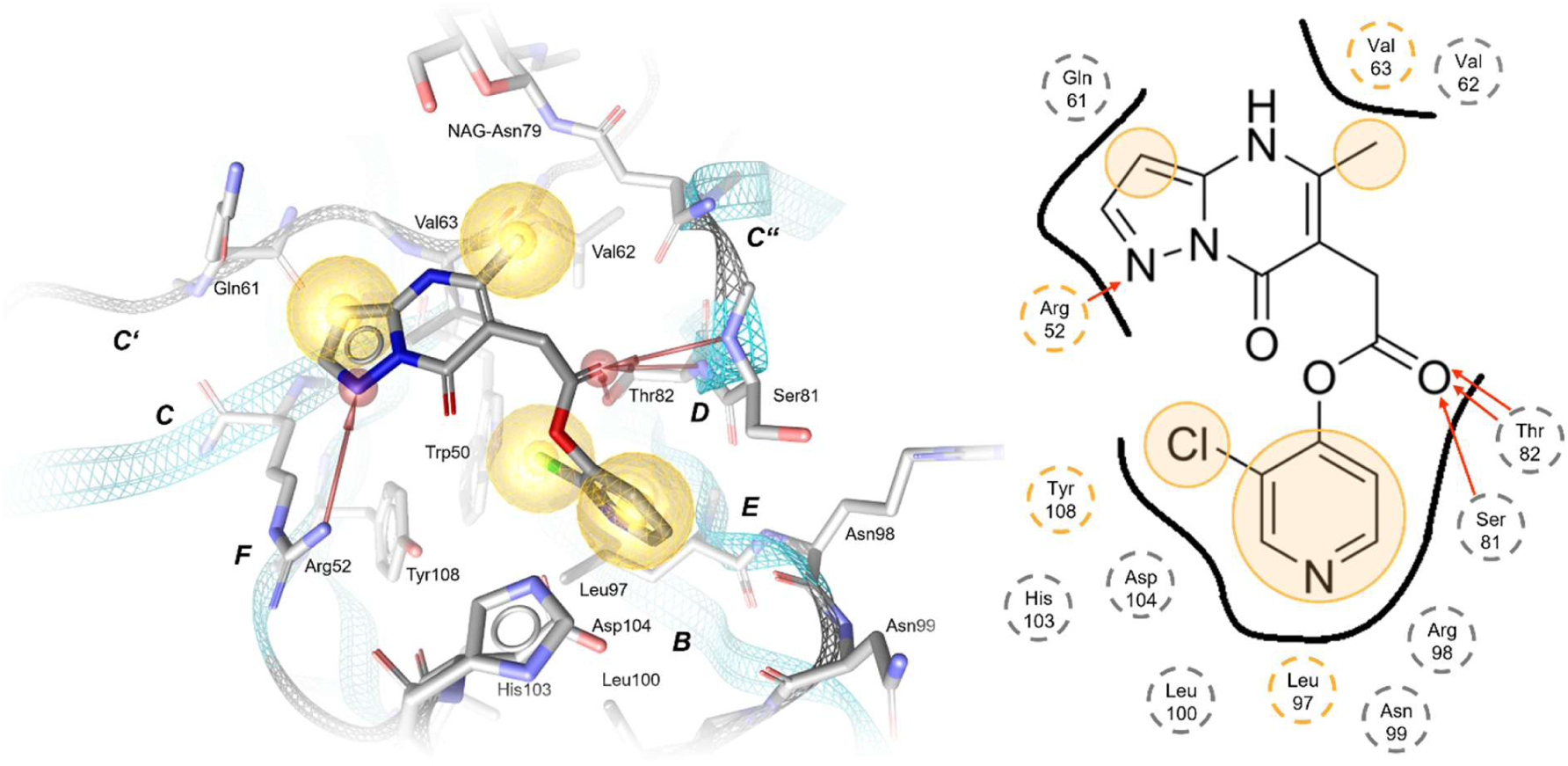
**Docking pose of EN020 with in TREM2 cavity with 3D pharmacophore**. Yellow/orange spheres depict hydrophobic contacts, and red arrows depict hydrogen bond acceptors.

### Preliminary Structure-Activity Relationship (SAR) study focused on the top hit compound, EN020

To further explore the SAR of the top hit compound **EN020**, a series of analogs was synthesized based on the pyrazolo[1,5-a]pyrimidinone core. The synthetic route commenced with 1*H*-pyrazol-5-amine (**1**), which underwent cyclization with dimethyl 2-acetylsuccinate (**2**) in xylene, furnishing compound **3** (**Scheme 1**). Ester hydrolysis of compound **3** yielded the corresponding acid (compound **4**), which was subsequently subjected to amide coupling with various substituted anilines to generate a set of analogs labeled **EN5–EN7**. These compounds feature a 7-oxo core and diverse aryl amide substituents, allowing exploration of electronic and steric effects. **EN5** bears a 4-chlorophenyl group, targeting potential halogen bonding and electron-withdrawing influence. **EN6** includes a 2,4-dimethylphenyl moiety, probing increased hydrophobicity and steric bulk. **EN7** combines a 3-chloro-4-methylphenyl ring to evaluate synergistic electron-donating and withdrawing effects on binding.

A second round of synthesis focused on aromatizing the core ring and introducing a piperidine moiety at the 7-position. The strategy involved chlorination of the core scaffold using phosphorus oxychloride (POCl₃), followed by nucleophilic substitution with dimethylamine, giving compound **8**. Subsequently, the chloro intermediate underwent nucleophilic aromatic substitution (SNAr) with piperidine in DMF, affording compound **9**. Basic hydrolysis with NaOH provided the corresponding acid (compound **10**), which served as a key intermediate for diversification. Using compound **10** as a central scaffold, a focused set of amide and ester derivatives (**ENP11–ENP13**) was synthesized via EDCI-mediated coupling with various alcohols and amines (**Scheme 1**). These modifications were designed to fine-tune properties such as hydrophobicity, hydrogen bonding capacity, and electronic character at the terminal group. **ENP-11** features a 2-chloropyridin-3-yl amide, integrating both heteroaromaticity and basic nitrogen to promote potential π-stacking and H-bonding. **ENP-12** and **ENP-13** are ester analogs incorporating chloropyridinyl alcohols, differing in substitution pattern (2-chloropyridin-3-yl vs. 3-chloropyridin-2-yl), to assess how regioisomerism affects binding orientation, hydrolytic stability, and target interaction.

**Scheme 1.**
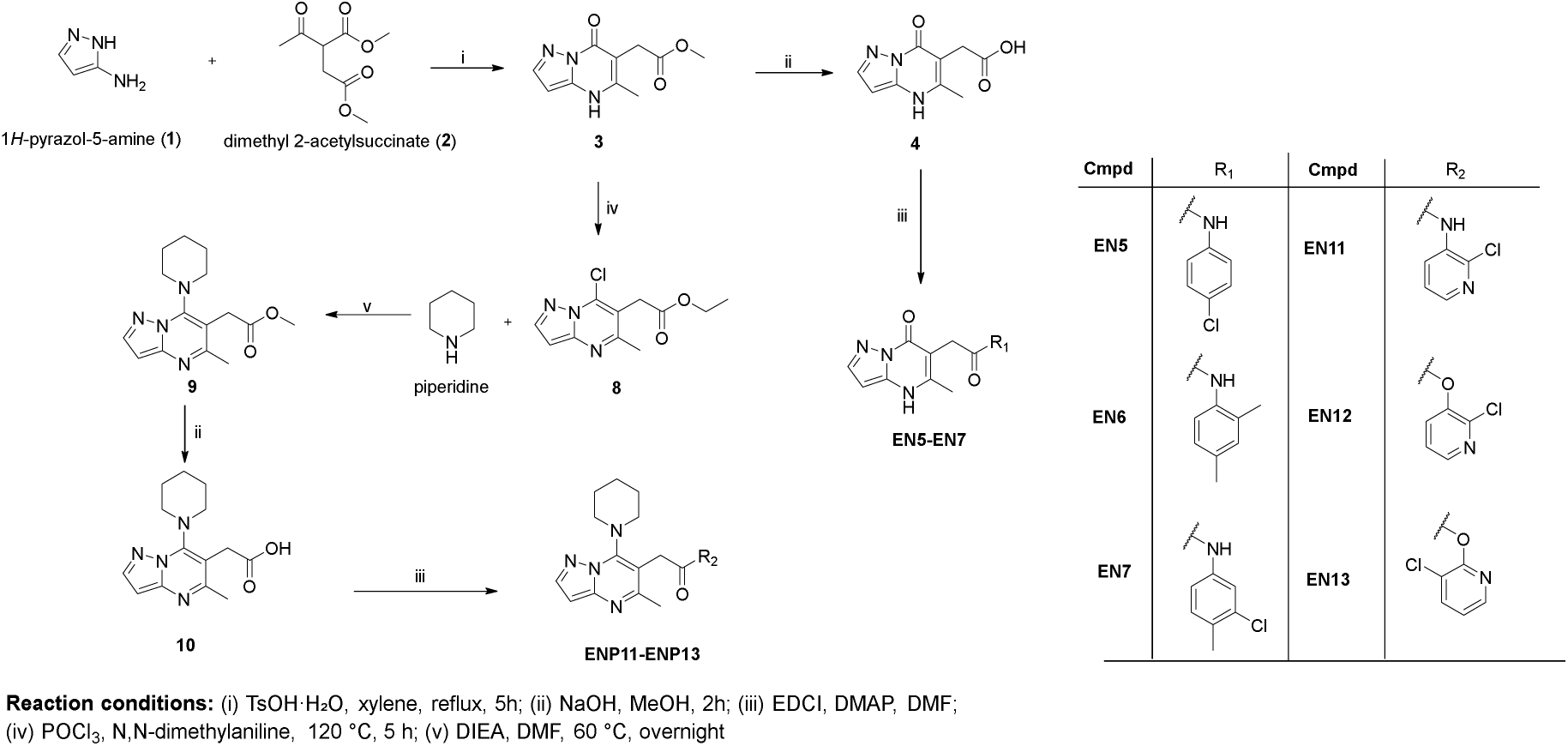
Synthesis of structural derivatives of the top hit compound EN020.

We complemented our synthetic campaign with a limited SAR-by-catalog strategy by sourcing two structural analogs of EN020, designated **T2V001** and **T2V002** (**Table 1**). A summary of the TREM2 binding affinities of **EN020** derivatives is presented in **Table 1**. Among the investigated compounds, **T2V002**, which incorporates a bulky cyclooctane moiety, retained measurable binding affinity (KD = 73.6 ± 19.8 µM) but did not surpass **EN020** (KD = 14.2 ± 2.2 µM). The preservation of activity in **T2V002** despite reduced potency suggests that lipophilic bulk can be accommodated in the binding pocket, possibly engaging peripheral hydrophobic contacts, but may impose conformational constraints that weaken key interactions responsible for high-affinity binding. In contrast, most other analogs, including those featuring halogenated aromatics, heteroaryl amides, and ester substitutions, showed negligible binding by MST. These outcomes point to a narrow tolerance for scaffold modification, where even minor electronic or steric changes at critical positions can disrupt the delicate balance of hydrophobic contacts and hydrogen-bonding interactions observed in the original **EN020** docking model.

**Table 1.**
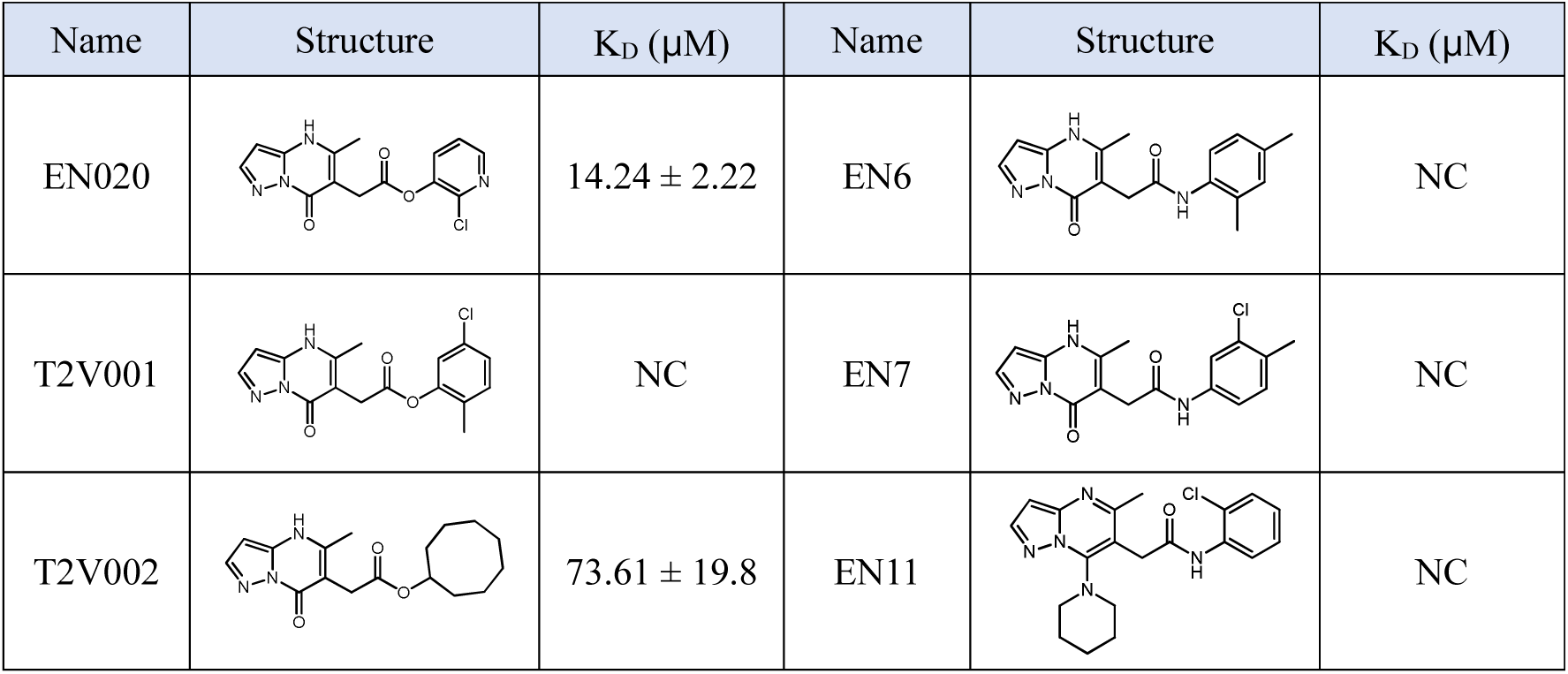
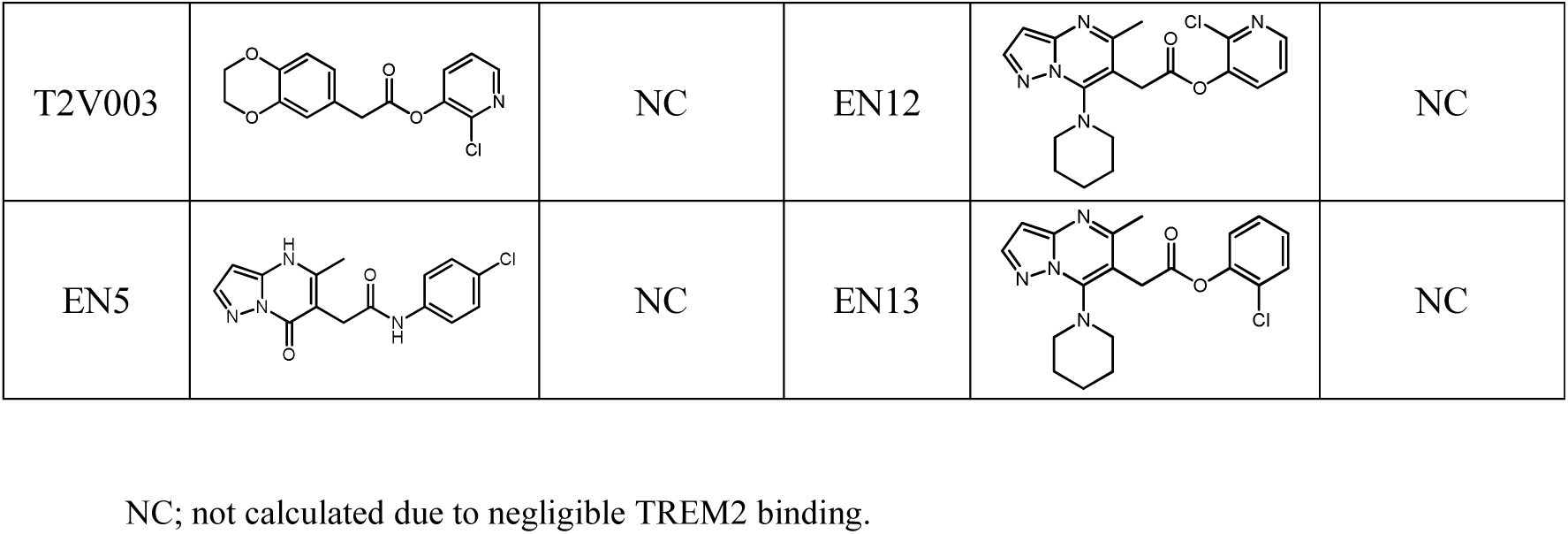
Chemical structures of EN020 derivatives and their corresponding TREM2 binding affinity.

| Name | Structure | $K_D$ ($\mu\text{M}$) | Name | Structure | $K_D$ ($\mu\text{M}$) |
| --- | --- | --- | --- | --- | --- |
| EN020 | | $14.24 \pm 2.22$ | EN6 | | NC |
| T2V001 |  | NC | EN7 |  | NC |
| T2V002 | | $73.61 \pm 19.8$ | EN11 | | NC |
| T2V003 |  | NC | EN12 |  | NC |
| EN5 |  | NC | EN13 |  | NC |

NC; not calculated due to negligible TREM2 binding.

Overall, this SAR exploration highlights **EN020** as the most potent TREM2-binding hit identified to date in our study, while **T2V002** serves as a proof-of-concept that selective scaffold modifications can maintain partial activity. These insights provide a starting point for rational optimization efforts aimed at preserving the core pharmacophoric features of **EN020** while fine-tuning physicochemical properties for improved potency, stability, and CNS penetration.

## Conclusion

Through an integrated structure-based virtual screening and experimental validation workflow, we identified **EN020** as a novel small molecule modulator of TREM2 that engages a putative allosteric site, binds with micromolar affinity, and enhances microglial phagocytic activity beyond that of a benchmark TREM2 agonist. Our SAR exploration revealed that the **EN020**–TREM2 interaction is highly sensitive to structural changes, with only limited modifications preserving activity, as exemplified by **T2V002**. These findings not only establish **EN020** as a promising lead for further development but also provide valuable pharmacophore and SAR insights to guide next-generation TREM2 modulators. Future efforts will focus on potency optimization, pharmacokinetic profiling, and in vivo efficacy studies to assess therapeutic potential in models of Alzheimer’s disease and related neurodegenerative disorders.

## Methods

### Structure preparation

The crystal structure of the extracellular Ig-like domain of human TREM2 (PDB: 5UD7)^[26]^ was used as a template for the entire workflow. To generate the template, the raw PDB structure was prepared in MOE (v2022.02 Chemical Computing Group ULC, 910-1010 Sherbrooke St. W., Montreal, QC H3A 2R7, 2024). Of the six chains in the crystal bundle, chain A was selected, due to its high resolution in the PDB file. The chain ends were then capped, and hydrogens were added by employing the Protonate3D algorithm^[37]^. Ligandscout’s (v4.4.3)^[38]^ built-in pocket detection was utilized to identify possible cavities eligible for targeting.

An additional full model of the full TREM2 was generated using Robetta’s structure prediction tool.^[39]^ The full TREM2 sequence was retrieved from UniProt (Q9NZC2). The structure of the extracellular Ig-like domain (PDB: 5UD7)^[26]^, and the transmembrane domain (PDB: 6Z0G)^[40]^ were also provided as templates for structure generation. The model that matched the two templates best was selected for further processing. It was then loaded into MOE (v2022.02, Chemical Computing Group ULC, 910–1010 Sherbrooke St. W., Montreal, QC, H3A 2R7, Canada, 2024). Missing atoms were added and the protein was protonated using Protonate3D^[37]^. Finally, Ramachandran outliers and atom clashes were minimized using the AMBER:ETH force field in MOE(v2022.02 Chemical Computing Group ULC, 910-1010 Sherbrooke St. W., Montreal, QC H3A 2R7, 2024).

### Molecular dynamics (MD) simulation and PyRod pharmacophore

The template was prepared for MD simulation in Maestro (v13.1.137; Schroedinger, LLC, New York City, NY, USA) first, by placing the structure in a box with a minimum buffer of 10 Å between the edges of the box and the protein. The box was then filled with 0.15 M sodium chloride dissolved in TIP4P water^[41]^. MD simulations were then performed ten times with the resulting system in Desmond (v2022.01; Schroedinger, LLC, New York City, NY, USA)^[42]^ for 10 ns, recording 1 frame every 5 ps, on NVIDIA GeForce GTX 2090 graphics cards using the OPLS force field^[43]^, resulting in ten replicas with 2000 frames each. The temperature of the system was maintained at 300 K employing the Nosé-Hoover thermostat^[44]^, while the pressure was maintained at 1.01325 bar using the Martyna-Tobias-Klein barostat^[45]^. VMD^[46]^ (v.1.9.3; University of Illinois Champaign, IL, USA) was used to post-process the MD simulations. The last thousand frames of each replica were then submitted to PyRod^[25]^ to generate dynamic molecular interaction fields (dMIFs) in the selected cavity. Pharmacophore features were then placed in the dMIFs (thresholds: HA_combo = 26 kcal/mol; HD_combo = 34 kcal/mol; HI = 87 kcal/mol) with Ligandscout resulting in one preliminary pharmacophore.

### Pharmacophore refinement and screening

Refinement of the preliminary pharmacophore and pharmacophore screening were conducted in Ligandscout. The ESC was screened against the preliminary pharmacophores in multiple cycles. In each cycle, pharmacophore features were added and omitted and adjusted in position, to improve enrichment of the hits and their binding modes and their shape complimentarity with the binding cavity, which resulted in one core pharmacophore with three optional pharmacophore features.

### Molecular Docking

The binding poses of the 1312 hits were validated by performing molecular docking in GOLD (v5.8.1; Genetic Optimization for Ligand Docking, The Cambridge Crystallographic Data Center, UK).^[47]^ First, each compound was enumerated in MOE at pH 7.4 to account for each protonation state of every individual compound. For each protonation state, ten binding hypotheses were built in the selected binding pocket. Atoms in a radius of 10 Å around X = 40.57, Y = -0.06 and Z = 8.58, which is located close to Trp50, form the binding site for the molecular docking step. During the molecular docking step, a rotamer of Arg52 was used, to facilitate the aforementioned hydrogen acceptor bond to Arg52. The rotamer used has the lowest potential energy in the Amber10 force field.^[48]^ Binding hypotheses with less than three hydrophobic features were discarded. The resulting 9540 binding hypotheses were then visually inspected.

### Allosteric Path Analysis

The MD simulation inputs for allosteric path analysis were prepared using CharmmGUI.^[49,50]^ The full TREM2 model was protonated at pH 7.4. It was then embedded in a POPC lipid bilayer aligned along a vector between the residues Ile175 and Ala195.^[51]^ The system was solvated in a TIP3P water box containing 0.15M NaCl. Simulations were performed using Charmm36m^[52]^ force fields and run in OpenMM^[53]^. Pre-production minimization and equilibration were performed according to the CharmmGUI protocol.^[49,50]^ Subsequently, three production replicas were executed for 200 ns (600 ns in total) in an NPγT ensemble at 303.15 K, 1.0 bar, and a surface tension of 0 dyne/cm. Allosteric path analysis was conducted by running MDPath^[54]^ with base settings on each replica seperatly.

### Temperature-Related Intensity Change (TRIC) Analysis

TRIC-based binding interactions were assessed utilizing a Dianthus NT.23PicoDuo system (NanoTemper Technologies, Munich, Germany). Human recombinant TREM2 protein (BioTechne, Minneapolis, MN, USA) underwent fluorescent labeling with RED-tris-NTA 2nd Generation dye through the His-Tag Labeling Kit (NanoTemper Technologies), following standard manufacturer guidelines. Binding experiments were carried out in PBST buffer (154 mM NaCl, 5.6 mM Na₂HPO₄, 1.05 mM KH₂PO₄, pH 7.4, supplemented with 0.005% Tween- 20). Fluorescently-labeled TREM2 was maintained at 10 nM final concentration for all measurements. Test substances underwent serial dilution in PBST buffer and were pre-incubated with labeled TREM2 protein for 10 minutes at ambient temperature (22-25°C) before analysis. Instrument settings included 85% LED excitation power, deactivated picomolar detector sensitivity, and 5-second laser exposure duration. Data processing involved initial analysis through Dianthus Analysis software (NanoTemper Technologies), with subsequent curve fitting and statistical evaluation performed using GraphPad Prism 10.0 (GraphPad Software, San Diego, CA, USA). PC-192 (100 μM), a recognized TREM2 agonist, served as the positive control.^[55]^

### Microscale Thermophoresis (MST) Analysis

Binding kinetics were evaluated through microscale thermophoresis using a Monolith NT.115 instrument (NanoTemper Technologies, Munich, Germany). Human recombinant TREM2 protein (BioTechne, Minneapolis, MN, USA) was fluorescently tagged using the RED-tris-NTA His-tag labeling system (NanoTemper Technologies) per manufacturer specifications. MST analyses were executed in PBS buffer (pH 7.4) supplemented with 0.005% Tween-20. Fluorescent-labeled proteins were employed at 40 nM final concentration and subjected to 10-minute room temperature (22-25°C) incubation with serially diluted test compounds before measurement. MST analysis utilized standard capillaries under the following conditions: red filter configuration, 100% LED intensity, and medium MST power setting. Thermophoretic movement was tracked for 20 seconds with an additional 5-second pre-measurement delay. Initial data processing was performed using MO.Affinity Analysis software (NanoTemper Technologies), with subsequent curve analysis conducted via GraphPad Prism 10.0 (GraphPad Software, San Diego, CA, USA).

### Surface Plasmon Resonance (SPR)

Binding of EN020 to human TREM2 protein was assessed using surface plasmon resonance (SPR) on a Biacore 8K system (Cytiva). Biotinylated TREM2 (Cat. No. 11084- H49H-B, Sino Biological) was immobilized on an SA Sensor Chip (Cytiva) in PBS-P buffer (0.2 M phosphate buffer, 27 mM KCl, 1.37 M NaCl, 0.5% Surfactant P20, pH 7.4) to a level of 4,343 ± 399 RU. Prior to immobilization, the chip surface was conditioned with three 1- minute injections of 1 M NaCl in 50 mM NaOH. After immobilization, the surface was washed sequentially with 50% (v/v) isopropanol in water, 1 M NaCl, and 50 mM NaOH to remove weakly or non-specifically bound material, then equilibrated in running buffer until a stable baseline was reached. Kinetic measurements were performed using a single-cycle kinetic method. Enamine 20 was tested at seven concentrations (200, 100, 50, 25, 12.5, 6.25 and 3.125 μM) in PBS-P containing 2% DMSO (Cytiva) and injected over the immobilized protein at 25°C, with a flow rate of 30 μL/min, contact time of 200 s, and dissociation time of 900 s. After each cycle, the surface was washed with 50% DMSO. The result is presented as sensorgram obtained after subtraction of the background response signal from a reference flow cell and from a control experiment with buffer injection. Interaction was investigated at least in triplicate. Data were analyzed using Biacore™ Insight Evaluation Software (Cytiva), with a 1:1 binding model providing the best fit to the experimental data.

### Phagocytosis Assay

Phagocytosis assay was performed in BV2 cells using greenfluorescent latex beads (Sigma #L1030). Prior use, beads were opsonized in FBS (1 h at 37 °C), the final concentrations for beads and FBS in DMEM were 0.01% (v/v) and 0.05% (v/v) respectively. Briefly, BV2 cells were submitted to serum starvation prior treatment with either compound **VG-3928** and **EN020** (25 μM for 1 h) or solubilization buffer (control condition) and beads were added to the media for an additional 30 min. Cultures were then washed 3 times with ice-cold PBS and fixed in 4% paraformaldehyde (PFA) (Thermofisher, MA, USA) before undergoing immunocytochemistry. Following Immunostaining, pictures were taken for each condition under 20X objective, total number of cells was counted manually using DAPI staining and each IBA1 immunolabelled-cell containing at least one bead was counted as positive for phagocytosis.

## Data Availability Statement

The data that support the findings of this study are available in the Supporting Information.

## Acknowledgement

This work was supported by the National Institute on Aging under grant number R01AG083512 (PI: Gabr).

## References

[1] M. Revi, Alzheimer’s Disease Therapeutic Approaches. Adv Exp Med Biol. 2020, 1195, 105–116.

[2] M. Tolar, S. Abushakra, M. Sabbagh, The path forward in Alzheimer’s disease therapeutics: Reevaluating the amyloid cascade hypothesis. Alzheimers Dement. 2020, 16 (11),1553–1560.

[3] R. Ismail, P. Parbo, L. Madsen, A. Hansen, J. Schaldemose, P. Kjeldsen, M. Stokholm, H. Gottrup, S. Eskildsen, D. Brooks, The relationships between neuroinflammation, beta-amyloid and tau deposition in Alzheimer’s disease: a longitudinal PET study. J Neuroinflammation. 2020, 17 (1), 151.

[4] D. Garbuz, O. Zatsepina, M. Evgen’ev, Beta Amyloid, Tau Protein, and Neuroinflammation: An Attempt to Integrate Different Hypotheses of Alzheimer’s Disease Pathogenesis. Mol Biol. 2021, 55 (5), 734–747.

[5] P. Cisternas, X. Taylor, C. Lasagna-Reeves, The Amyloid-Tau-Neuroinflammation Axis in the Context of Cerebral Amyloid Angiopathy. Int J Mol Sci. 2019, 20 (24), 6319.

[6] Q. Li, B. Barres, Microglia and macrophages in brain homeostasis and disease. Nat Rev Immunol. 2018, 18 (4), 225–242.

[7] W. Fu, X. Wang, N. Ip, Targeting Neuroinflammation as a Therapeutic Strategy for Alzheimer’s Disease: Mechanisms, Drug Candidates, and New Opportunities. ACS Chem Neurosci. 2019, 10 (2), 872–879.

[8] R. Dhapola, S. Hota, P. Sarma, A. Bhattacharyya, B. Medhi, D. Reddy, Recent advances in molecular pathways and therapeutic implications targeting neuroinflammation for Alzheimer’s disease. Inflammopharmacology. 2021, 29 (6), 1669–1681.

[9] A. Deczkowska, A. Weiner, I. Amit, The Physiology, Pathology, and Potential Therapeutic Applications of the TREM2 Signaling Pathway. Cell. 2020, 181 (6),1207–1217.

[10] S. Carmona, K. Zahs, E. Wu, K. Dakin, J. Bras, R. Guerreiro, The role of TREM2 in Alzheimer’s disease and other neurodegenerative disorders. Lancet Neurol. 2018, 17 (8), 721–730.

[11] M. Gratuze, C. Leyns, D. Holtzman, New insights into the role of TREM2 in Alzheimer’s disease. Mol Neurodegener. 2018, 13 (1), 66.

[12] T. Ulland, M. Colonna, TREM2 — a key player in microglial biology and Alzheimer disease. Nat Rev Neurol. 2018, 14 (11), 667–675.

[13] Y. Wang, M. Cella, K. Mallinson, J. Ulrich, K. Young, M. Robinette, S. Gilfillan, G. Krishnan, S. Sudhakar, B. Zinselmeyer, D. Holtzman, J. Cirrito, M. Colonna, TREM2 lipid sensing sustains the microglial response in an Alzheimer’s disease model. Cell. 2015, 160 (6), 1061–71.

[14] P. Yuan, C. Condello, C. Keene, Y. Wang, T. Bird, S. Paul, W. Luo, M. Colonna, D. Baddeley, J. Grutzendler, TREM2 Haplodeficiency in Mice and Humans Impairs the Microglia Barrier Function Leading to Decreased Amyloid Compaction and Severe Axonal Dystrophy. Neuron. 2016, 90 (4), 724–739.

[15] H. Keren-Shaul, A. Spinrad, A. Weiner, O. Matcovitch-Natan, R. Dvir-Szternfeld, T. Ulland, E. David, K. Baruch, D. Lara-Astaiso, B. Toth, S. Itzkovitz, M. Colonna, M. Schwartz, I. Amit, A Unique Microglia Type Associated with Restricting Development of Alzheimer’s Disease. Cell. 2017, 169 (7), 1276–1290.

[16] C. Lee, A. Daggett, X. Gu, L. Jiang, P. Langfelder, X. Li, N. Wang, Y. Zhao, C. Park, Y. Cooper, I. Ferando, I. Mody, G. Coppola, H. Xu, X. Yang, Elevated TREM2 Gene Dosage Reprograms Microglia Responsivity and Ameliorates Pathological Phenotypes in Alzheimer’s Disease Models. Neuron. 2018, 97 (5),1032–1048.

[17] K. Schlepckow, K. Monroe, G. Kleinberger, L. Cantuti-Castelvetri, S. Parhizkar, D. Xia, M. Willem, G. Werner, N. Pettkus, B. Brunner, A. Sülzen, B. Nuscher, H. Hampel, X. Xiang, R. Feederle, S. Tahirovic, J. Park, R. Prorok, C. Mahon, C. Liang, J. Shi, D. Kim, H. Sabelström, F. Huang, G. Di Paolo, M. Simons, J. Lewcock, C. Haass, Enhancing protective microglial activities with a dual function TREM2 antibody to the stalk region. EMBO Mol Med. 2020, 12(4), e11227.

[18] B. Price, T. Sudduth, E. Weekman, S. Johnson, D. Hawthorne, A. Woolums, D. Wilcock, Therapeutic Trem2 activation ameliorates amyloid-beta deposition and improves cognition in the 5XFAD model of amyloid deposition. J Neuroinflammation. 2020, 17 (1), 238.

[19] M. Cavaco, D. Gaspar, M. Arb Castanho, V. Neves, Antibodies for the Treatment of Brain Metastases, a Dream or a Reality? Pharmaceutics. 2020, 12 (1), 62.

[20] L. A. Lampson, Monoclonal antibodies in neuro-oncology: Getting past the blood-brain barrier. MAbs. 2011, 3(2), 153–160.

[21] W. M. Pardridge, The blood-brain barrier: bottleneck in brain drug development. NeuroRx. 2005, 2 (1), 3–14.

[22] E. A. Chowdhury, B. Noorani, F. Alqahtani, A. Bhalerao, S. Raut, F. Sivandzade, L. Cucullo, Understanding the brain uptake and permeability of small molecules through the BBB: A technical overview. J Cereb Blood Flow Metab. 2021, 41(8), 1797–1820.

[23] M. Gabr, A. U. Rehman, H. Chen, Quinoline-Pyrazole Scaffold as a Novel Ligand of Galectin-3 and Suppressor of TREM2 Signaling. ACS Med Chem Lett. 2020, 11 (9), 1759–1765.

[24] C. Mirescu, Characterization of the first TREM2 small molecule agonist, VG-3927, for clinical development in Alzheimer’s disease. Alzheimers Dement. 2025, *20* (Suppl 6), e084622.

[25] D. Schaller, S. Pach. G. Wolber, PyRod: Tracing Water Molecules in Molecular Dynamics Simulations. J Chem Inf Model. 2019, 59(6), 2818–2829.

[26] A. Sudom, S. Talreja, J. Danao, E. Bragg, R. Kegel, X. Min, J. Richardson, Z. Zhang, N. Sharkov, E. Marcora, S. Thibault, J. Bradley, S. Wood, A. Lim, H. Chen, S. Wang, I. Foltz, S. Sambashivan, Z. Wang, Molecular basis for the loss-of-function effects of the Alzheimer’s disease-associated R47H variant of the immune receptor TREM2. J Biol Chem 2018, 293(32), 12634–12646.

[27] G. Wolber, T. Langer, LigandScout: 3-D pharmacophores derived from protein-bound ligands and their use as virtual screening filters. J Chem Inf Model 2005, 45(1), 160–169.

[28] J. Abascal, C. Vega, A general purpose model for the condensed phases of water: TIP4P/2005. Journal of Chemical Physics, 2005, 123(23).

[29] K. J. Bowers, et al., Scalable algorithms for molecular dynamics simulations on commodity clusters, in Proceedings of the 2006 ACM/IEEE conference on Supercomputing. 2006, Association for Computing Machinery: Tampa, Florida. p. 84–es.

[30] W. L. Jorgensen, D. S. Maxwell, J. TiradoRives, Development and testing of the OPLS all atom force field on conformational energetics and properties of organic liquids. Journal of the American Chemical Society, 1996, 118(45), 11225–11236.

[31] G. Martyna, M.L. Klein, M. Tuckerman, Nosé–Hoover chains: The canonical ensemble via continuous dynamics. The Journal of Chemical Physics, 1992, 97(4), 2635–2643.

[32] G. Martyna, D.J. Tobias, M.L. Klein, Constant pressure molecular dynamics algorithms. The Journal of Chemical Physics, 1994, 101(5), 4177–4189.

[33] W. Humphrey, A. Dalke, K. Schulten, VMD: visual molecular dynamics. J Mol Graph, 1996, 14(1), p. 33–8 27-8.

[34] R. Ferreira de Freitas, M. Schapira, A systematic analysis of atomic protein-ligand interactions in the PDB. Medchemcomm, 2017, 8(10), 1970–1981.

[35] G. Jones, et al., Development and validation of a genetic algorithm for flexible docking. J Mol Biol, 1997, 267(3), 727–48.

[36] D. A. Case, et al., The Amber biomolecular simulation programs. J Comput Chem, 2005. 26(16), 1668–1688.

[37] Labute, P., Protnate 3D Assignment of ionization states and hydrogen coordinates to macromolecular structures, Proteins-Structure Function and Bioinformatics, 2005 75(1), 187–205

[38] Wolber, G. and T. Langer, LigandScout: 3-D pharmacophores derived from protein-bound ligands and their use as virtual screening filters. J Chem Inf Model, 2005. 45(1), 160–169.

[39] Baek, M., et al., Accurate prediction of protein structures and interactions using a three-track neural network. Science, 2021. 373(6557), 871-876.

[40] Steiner, A., et al., Gamma-Secretase cleavage of the Alzheimer risk factor TREM2 is determined by its intrinsic structural dynamics. EMBO J, 2020. 39(20), e104247.

[41] Abascal, J.L. and C. Vega, A general purpose model for the condensed phases of water: TIP4P/2005. J Chem Phys, 2005. 123(23), 234505.

[42] Bowers, K.J., et al., Scalable algorithms for molecular dynamics simulations on commodity clusters, in Proceedings of the 2006 ACM/IEEE conference on Supercomputing. 2006, Association for Computing Machinery: Tampa, Florida. p. 84–es.

[43] Jorgensen, W.L., D.S. Maxwell, and J. TiradoRives, Development and testing of the OPLS all-atom force field on conformational energetics and properties of organic liquids. J Am Chem Soc, 1996. 118(45), 11225–11236.

[44] Martyna, G.J., M.L. Klein, and M. Tuckerman, Nose-Hoover Chains - the Canonical Ensemble Via Continuous Dynamics. J Chem Phys, 1992. 97(4), 2635–2643.

[45] Martyna, G.J., D.J. Tobias, and M.L. Klein, Constant-Pressure Molecular-Dynamics Algorithms. J Chem Phys, 1994. 101(5), 4177–4189.

[46] Humphrey, W., A. Dalke, and K. Schulten, VMD: Visual molecular dynamics. J Mol Graph Model, 1996. 14(1), 33–38.

[47] Jones, G., et al., Development and validation of a genetic algorithm for flexible ligand docking. Abstracts of Papers of the American Chemical Society, 1997. 214: p. 154-Comp.

[48] Case, D.A., et al., The Amber biomolecular simulation programs. J Comput Chem, 2005. 26(16), 1668–1688.

[49] Jo, S., et al., CHARMM-GUI: A web-based graphical user interface for CHARMM. J Comput Chem, 2008. 29(11), 1859–1865.

[50] Brooks, B.R., et al., CHARMM: The biomolecular simulation program. J Comput Chem. 2009. 30(10), 1545–1614.

[51] Wu, E.L., et al., CHARMM-GUI Membrane Builder toward realistic biological membrane simulations. J Comput Chem. 2014. 35(27), 1997–2004.

[52] Huang, J., et al., CHARMM36m: an improved force field for folded and intrinsically disordered proteins. Nature Methods, 2017, 14(1), 71–73.

[53] Eastman, P., et al., OpenMM 8: Molecular Dynamics Simulation with Machine Learning Potentials. The Journal of Physical Chemistry B, 2024, 128(1), 109–116.

[54] Doering, N.P., et al., *MDPath: Unraveling Allosteric Communication Paths of Drug Targets through Molecular Dynamics Simulations*. 2025, Cold Spring Harbor Laboratory.

[55] Yuan, S., F. El Gaamouch, S. Cho, K. Kuncewicz, H. Nada and M. T. Gabr. ACS Med Chem Lett 2025, 16, 1634–1640.

