## Supporting Information for "Structure-Based Virtual Screening Identifies TREM2-Targeted Small Molecules that Enhance Microglial Phagocytosis"

*Electronic Supplementary Information*

| **Contents** |  |
| --- | --- |
| Chemistry | S2 |
| Chemical structures of hit compounds from virtual screening as potential TREM2 binders | S5 |
| NMR spectra of the synthesized compounds | S7 |
| Mass spectra of the synthesized compounds | S13 |

**Chemistry**

All reagents, solvents, and consumables were purchased from commercial suppliers and used as received, without further purification. ^1^H NMR spectra were recorded on a Bruker 500 MHz spectrometer. Chemical shifts (δ) are reported in parts per million (ppm) relative to the internal standard, with signal multiplicities designated as s, d, dd, t, q, td, dt, ddd, br, and m. Purification of compounds was conducted using flash chromatography on a CombiFlash system equipped with prepacked RediSep RF silica cartridges. Reaction progress and compound purity were monitored by thin-layer chromatography (TLC) on silica gel 60 F₂₅₄ plates.

**General Procedure for compounds 4 and 10:** To a solution of compound **3** or **9** in MeOH, an aqueous solution of NaOH (2.0 equiv) was added, and the mixture was stirred at room temperature for 2 h. Upon completion, MeOH was removed under reduced pressure, and the pH of the aqueous residue was adjusted to 2 using 1 M HCl, resulting in precipitation of the product. The solid was collected by filtration to afford desired compounds in quantitative yield.

**General Procedure for compounds** **EN5-7**and **EN11-13**: To a solution of the acid (100 mg, 1.0 equiv) in DMF (10 mL), the amine (1.2 equiv) was added, followed by DMAP (0.1 equiv). After stirring for 5 minutes, EDCI (1.5 equiv) was introduced, and the reaction mixture was stirred at room temperature for 24 hours. Upon completion, the reaction mixture was directly loaded onto a silica gel column and purified using a DCM/MeOH gradient to yield the desired compound.

**ethyl 2-(5-methyl-7-oxo-4,7-dihydropyrazolo[1,5-a]pyrimidin-6-yl)acetate** (**3**): A suspension of 1H-pyrazol-5-amine (**1**) (2.0 g, 24 mmol), diethyl 2-acetylsuccinate (4.69 mL, 28 mmol), and TsOH·H₂O (22.89 mg, 120 μmol) in o-xylene (125 mL) was refluxed using a Dean–Stark condenser for 5 h. Upon heating, the suspension became a clear homogeneous solution, and over time, a yellow solid began to precipitate from the mixture. After completion, the reaction mixture was cooled, diluted with hexanes (250 mL), filtered, washed with acetone, and dried to afford compound **3** as a light-yellow solid (90% yield).

**ethyl 2-(7-chloro-5-methylpyrazolo[1,5-a]pyrimidin-6-yl)acetate** (**8**): A solution of compound **3** (1.00 g, 4.25 mmol) and N,N-dimethylaniline (1.08 mL, 8.50 mmol) in POCl₃ (25 mL) was heated at 120 °C for 5 h. The reaction mixture was cooled to room temperature, concentrated *in vacuo*, and poured into a large quantity of ice-water. The resulting mixture was extracted with ethyl acetate (500 mL), and the organic layer was washed sequentially with water and brine (100 mL), dried over Na₂SO₄, filtered, and concentrated *in vacuo*. The crude product was triturated with ethyl acetate/hexane to afford **5** as a light-yellow solid (65% yield).

**ethyl 2-(5-methyl-7-(piperidin-1-yl)pyrazolo[1,5-a]pyrimidin-6-yl)acetate** (**9**): A mixture of compound **8** (1.00 g, 3.94 mmol), piperidine (0.428 mL, 4.34 mmol), and DIEA (0.428 mL, 4.34 mmol) in DMF (10 mL) was stirred at 60 °C overnight. Upon completion, the reaction mixture was cooled, diluted with diethyl ether (200 mL), and washed sequentially with water (3 × 50 mL) and brine (50 mL). The organic layer was dried over Na₂SO₄, filtered, and concentrated to afford a brown viscous oil. Purification by flash chromatography (hexane/ethyl acetate) yielded compound **6** as a yellow oil (90% yield).

**N-(4-chlorophenyl)-2-(5-methyl-7-oxo-4,7-dihydropyrazolo[1,5-a]pyrimidin-6-yl)acetamide (EN5):** white solid, 62% yield, ^1^H NMR (500 MHz, DMSO) δ 12.29 (s, 1H), 10.18 (s, 1H), 7.85 (d, *J* = 2.0 Hz, 1H), 7.63 (d, *J* = 8.9 Hz, 2H), 7.36 (d, *J* = 8.8 Hz, 2H), 6.11 (d, *J* = 1.9 Hz, 1H), 3.61 (s, 2H), 2.35 (s, 3H); ^13^C NMR (126 MHz, DMSO) δ 169.56, 157.43, 148.67, 143.39, 141.34, 138.71, 129.05, 127.05, 121.13, 100.73, 88.25, 33.23, 17.95; HRMS: calcd for C_15_H_14_N_4_O_2_Cl [M+H]^+^ m/z 317.0805, found 317.0807.

**N-(2,4-dimethylphenyl)-2-(5-methyl-7-oxo-4,7-dihydropyrazolo[1,5-a]pyrimidin-6-yl)acetamide (EN6):** white solid, 55% yield;^1^H NMR (500 MHz, DMSO) δ 12.28 (s, 1H), 9.24 (s, 1H), 7.85 (d, *J* = 1.9 Hz, 1H), 7.23 (d, *J* = 8.0 Hz, 1H), 7.02 (s, 1H), 6.95 (d, *J* = 8.2 Hz, 1H), 6.10 (d, *J* = 1.9 Hz, 1H), 3.59 (s, 2H), 2.37 (s, 3H), 2.24 (s, 3H), 2.16 (s, 3H);^13^C NMR (126 MHz, DMSO) δ 169.24, 157.56, 148.50, 143.33, 141.38, 134.59, 134.38, 132.24, 131.21, 126.83, 125.54, 101.00, 88.21, 40.49, 40.33, 40.16, 39.99, 39.82, 39.66, 39.49, 32.82, 20.94, 18.21, 17.95; HRMS: calcd for C_17_H_18_N_4_O_2_ [M+H]^+^ m/z 311.1508, found 311.1512.

**N-(3-chloro-4-methylphenyl)-2-(5-methyl-7-oxo-4,7-dihydropyrazolo[1,5-a]pyrimidin-6-yl)acetamide (EN7):** white solid, 62% yield; ^1^H NMR (500 MHz, DMSO) δ 12.30 (s, 1H), 10.13 (s, 1H), 7.85 (d, *J* = 2.0 Hz, 1H), 7.80 (d, *J* = 2.2 Hz, 1H), 7.38 (dd, *J* = 8.3, 2.2 Hz, 1H), 7.28 (d, *J* = 8.4 Hz, 1H), 6.11 (d, *J* = 2.1 Hz, 1H), 3.60 (s, 2H), 2.36 (s, 3H), 2.28 (s, 3H); ^13^C NMR (126 MHz, DMSO) δ 169.56, 157.43, 148.70, 143.38, 141.35, 138.88, 133.40, 131.60, 130.09, 119.58, 118.22, 100.69, 88.25, 33.22, 19.38, 17.96; HRMS: calcd for C_16_H_16_N_4_O_2_Cl [M+H]^+^ m/z 331.0962, found 331.0963.

**N-(2-chloropyridin-3-yl)-2-(5-methyl-7-(piperidin-1-yl)pyrazolo[1,5-a]pyrimidin-6-yl)acetamide (ENP11):** white solid, 50% yield, ^1^H NMR (500 MHz, CDCl_3_) δ 8.69 (dd, *J* = 8.1, 1.8 Hz, 1H), 8.13 (dd, *J* = 4.7, 1.7 Hz, 1H), 8.03 (d, *J* = 2.3 Hz, 1H), 7.77 (s, 1H), 7.30 – 7.26 (m, 1H), 6.56 (d, *J* = 2.3 Hz, 1H), 3.97 (s, 2H), 3.46 (s, 4H), 2.61 (s, 3H), 1.72 (h, *J* = 5.6 Hz, 6H); ^13^C NMR (126 MHz, CDCl_3_) δ 168.99, 159.76, 150.68, 149.41, 144.28, 144.03, 139.88, 131.51, 129.12, 123.46, 107.44, 95.63, 50.64, 36.72, 26.36, 23.94, 23.91; HRMS: calcd for C_19_H_21_N_6_OCl [M+H]^+^ m/z 385.1544, found 385.1544.

**2-chloropyridin-3-yl 2-(5-methyl-7-(piperidin-1-yl)pyrazolo[1,5-a]pyrimidin-6-yl)acetate (ENP12):** white solid, 50% yield; ^1^H NMR (500 MHz, DMSO) δ 8.39 (dd, *J* = 4.7, 1.7 Hz, 1H), 8.13 (d, *J* = 2.2 Hz, 1H), 7.94 (dd, *J* = 8.0, 1.6 Hz, 1H), 7.58 (dd, *J* = 8.0, 4.7 Hz, 1H), 6.54 (d, *J* = 2.2 Hz, 1H), 4.30 (s, 2H), 3.47 – 3.42 (m, 4H), 2.58 (s, 3H), 1.70 (dq, *J* = 22.7, 4.2 Hz, 6H); ^13^C NMR (126 MHz, DMSO) δ 169.74, 160.41, 150.32, 148.66, 147.85, 144.39, 143.67, 143.54, 133.79, 125.21, 106.67, 95.07, 50.48, 40.29, 40.13, 39.96, 39.79, 39.63, 26.36, 24.10, 23.67; HRMS: calcd for C_19_H_20_N_5_O_2_Cl [M+H]^+^ m/z 386.1384, found 386.1387.

**3-chloropyridin-2-yl 2-(5-methyl-7-(piperidin-1-yl)pyrazolo[1,5-a]pyrimidin-6-yl)acetate (ENP13):** white solid, 55% yield; ^1^H NMR (500 MHz, DMSO) δ 8.40 (dd, *J* = 4.8, 1.6 Hz, 1H), 8.18 (dd, *J* = 7.9, 1.6 Hz, 1H), 8.10 (d, *J* = 2.2 Hz, 1H), 7.48 (dd, *J* = 7.9, 4.8 Hz, 1H), 6.52 (d, *J* = 2.3 Hz, 1H), 4.29 (s, 2H), 3.41 (s, 4H), 2.56 (s, 3H), 1.75 – 1.62 (m, 6H); ^13^C NMR (126 MHz, DMSO) δ 168.55, 159.39, 152.60, 148.98, 148.16, 146.54, 143.15, 139.82, 124.08, 122.13, 94.26, 49.24, 32.12, 25.27, 23.07, 22.77; HRMS: calcd for C_19_H_20_N_5_O_2_Cl [M+H]^+^ m/z 386.1384, found 386.1385.

**Table S1.** Chemical structures of hit compounds from virtual screening as potential TREM2 binders.

| No. | Structure | No. | Structure |
| --- | --- | --- | --- |
| EN001 | 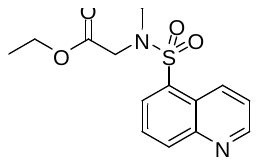 | EN011 | 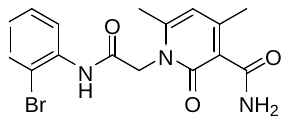 |
| EN002 | 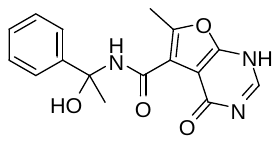 | EN012 | 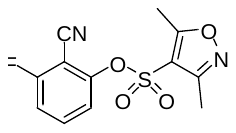 |
| EN003 | 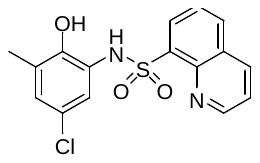 | EN013 | 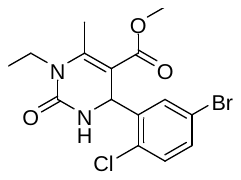 |
| EN004 | 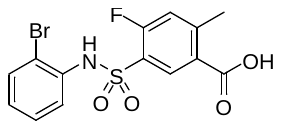 | EN014 | 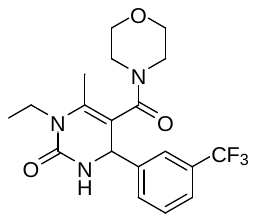 |
| EN005 | 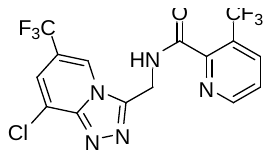 | EN015 | 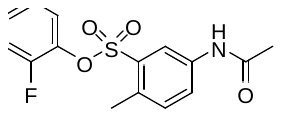 |
| EN006 | 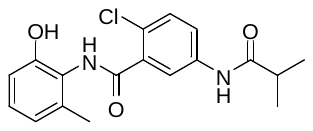 | EN016 | 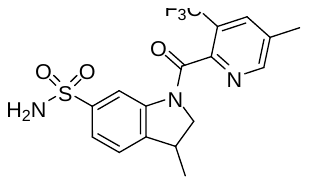 |
| EN007 | 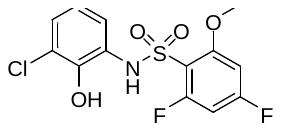 | EN017 | 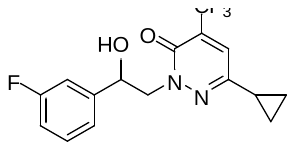 |
| EN008 | 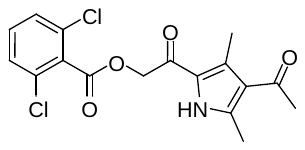 | EN018 | 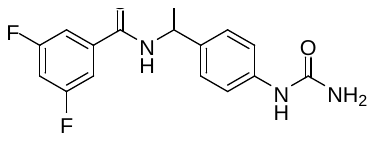 |
| EN009 | 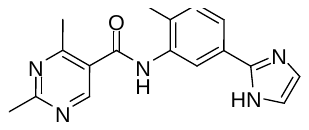 | EN019 | 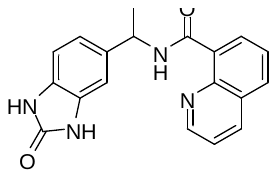 |
| EN010 | 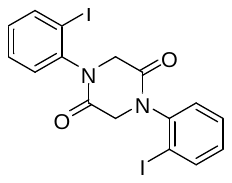 | EN020 | 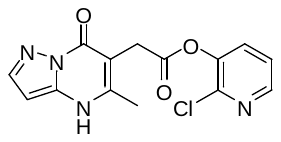 |

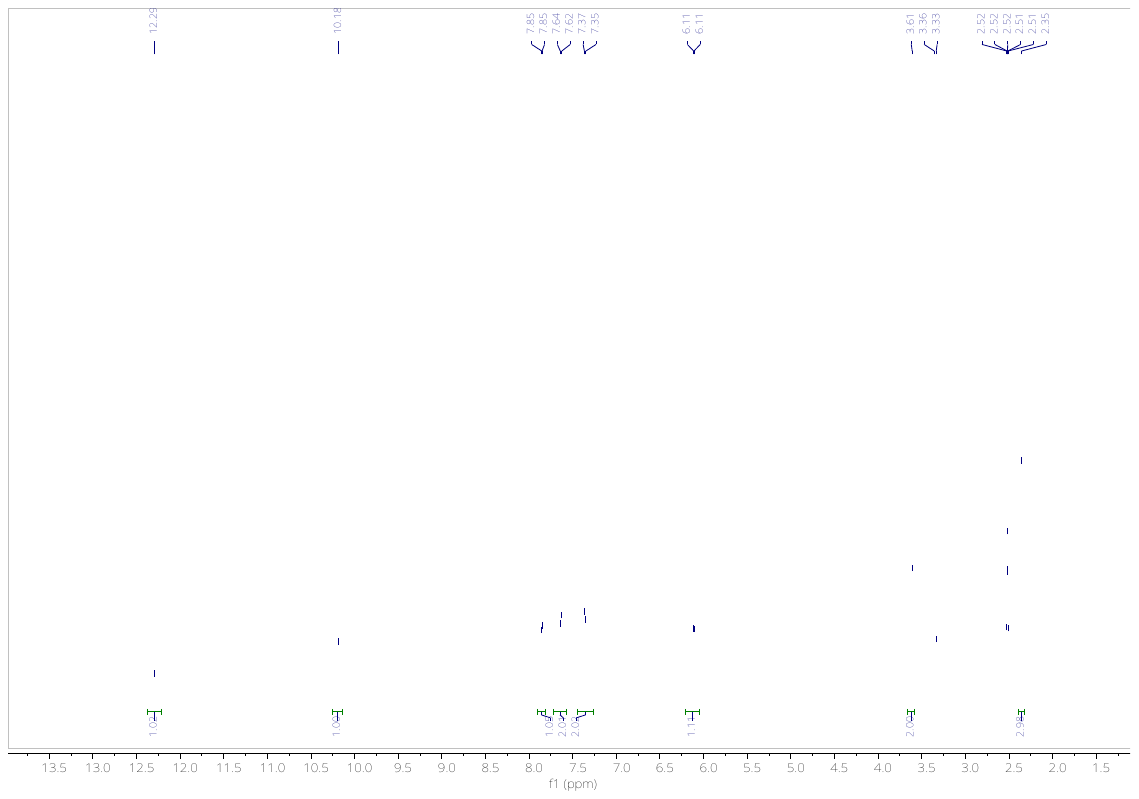

**Fig. 1.** ^1^H NMR of **EN5**.

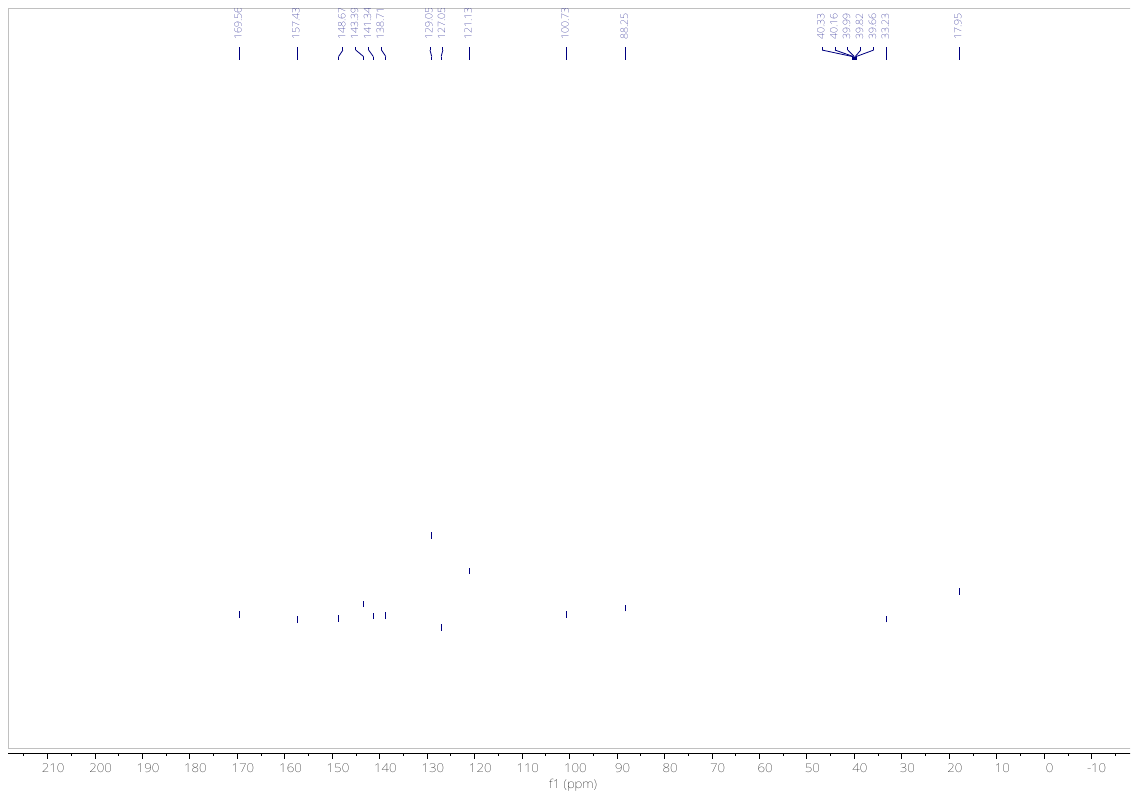

**Fig. 2.** ^13^C NMR of **EN5**.

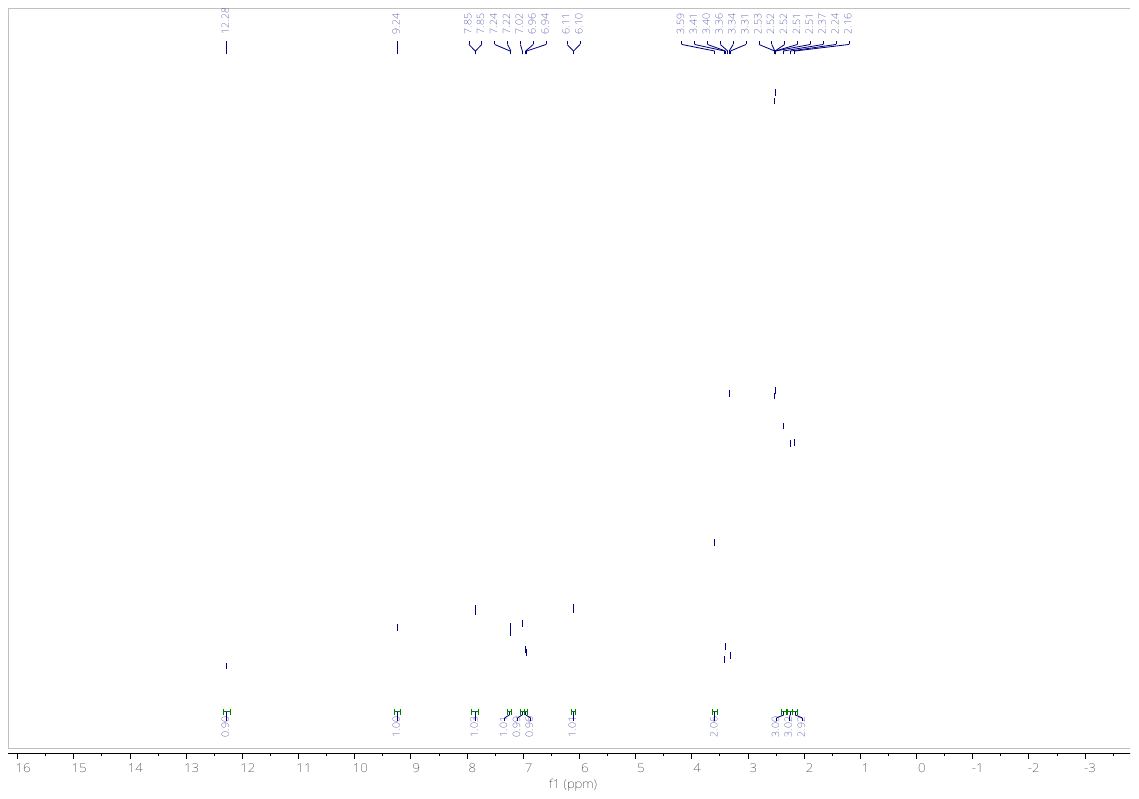

**Fig. 3.** ^1^H NMR of **EN6**.

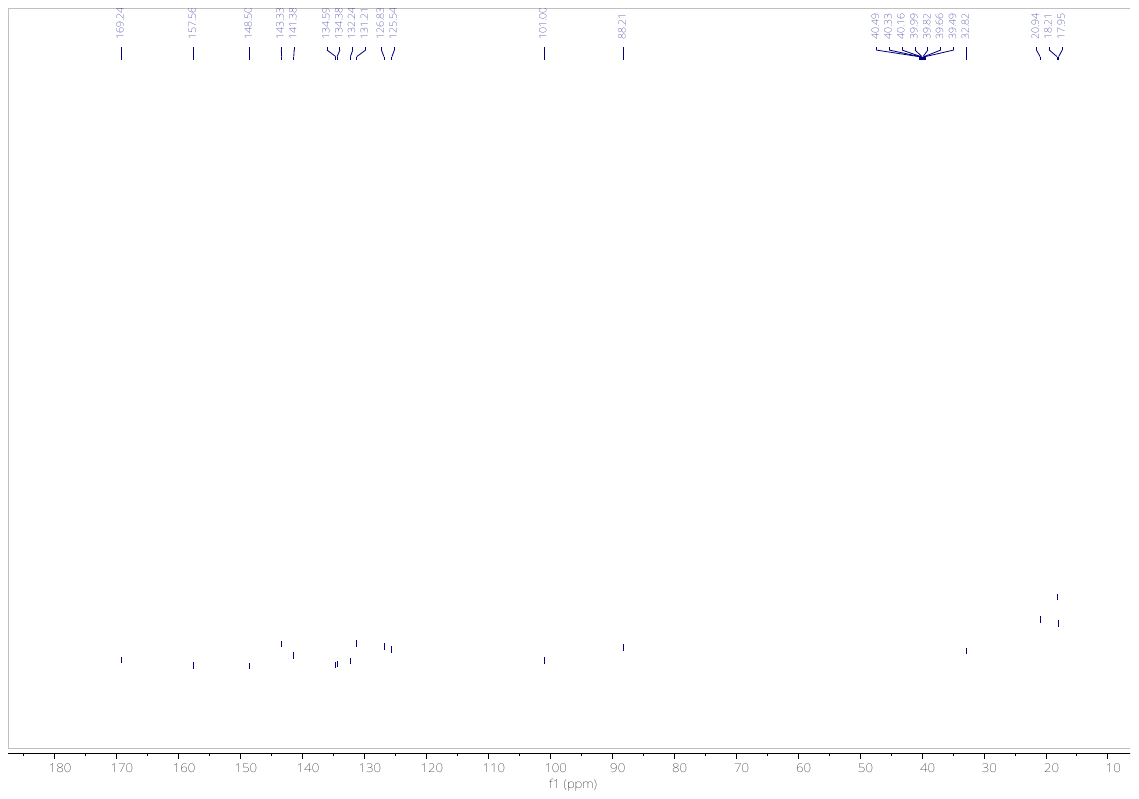

**Fig. 4.** ^13^C NMR of **EN6**.

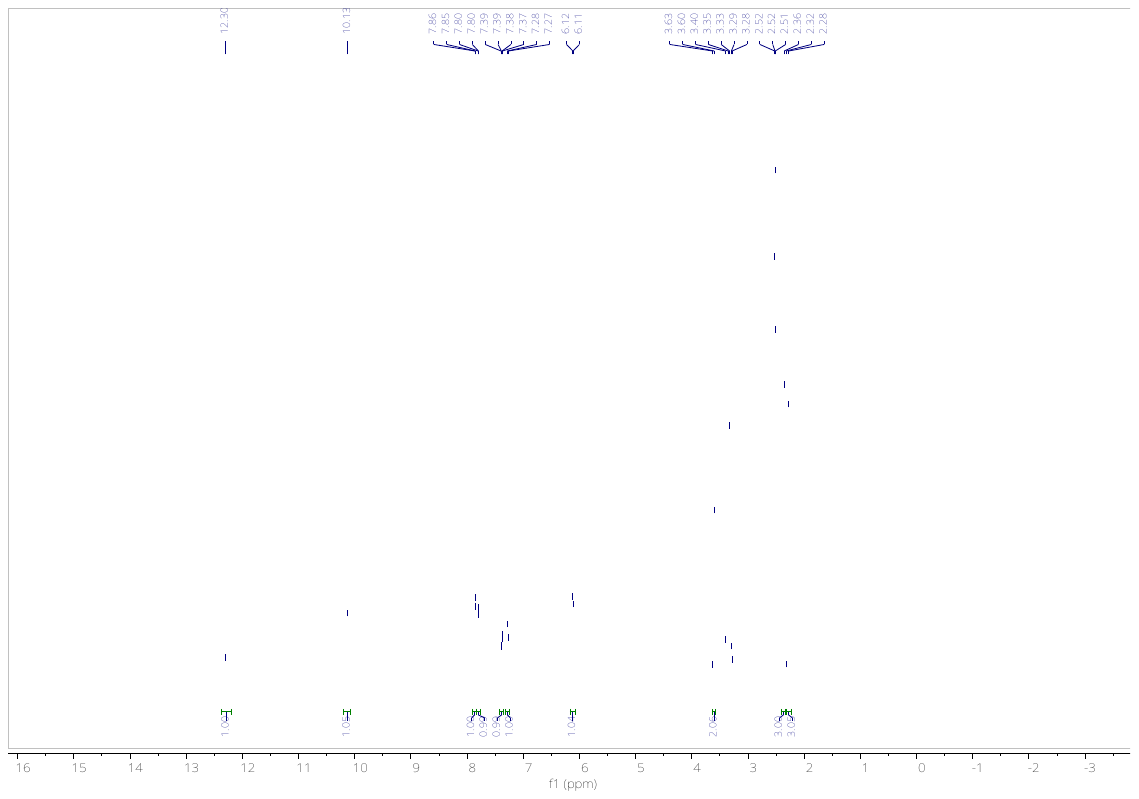

**Fig. 5.** ^1^H NMR of **EN7**.

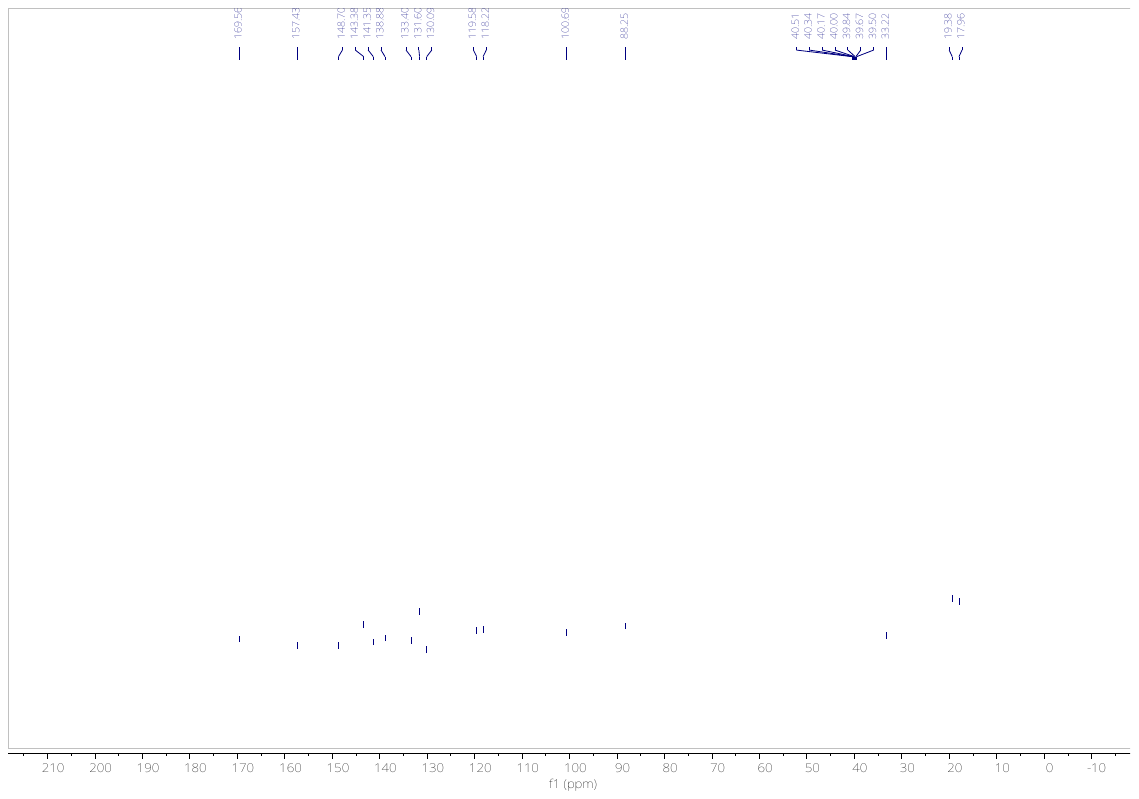

**Fig. 6.** ^13^C NMR of **EN7**.

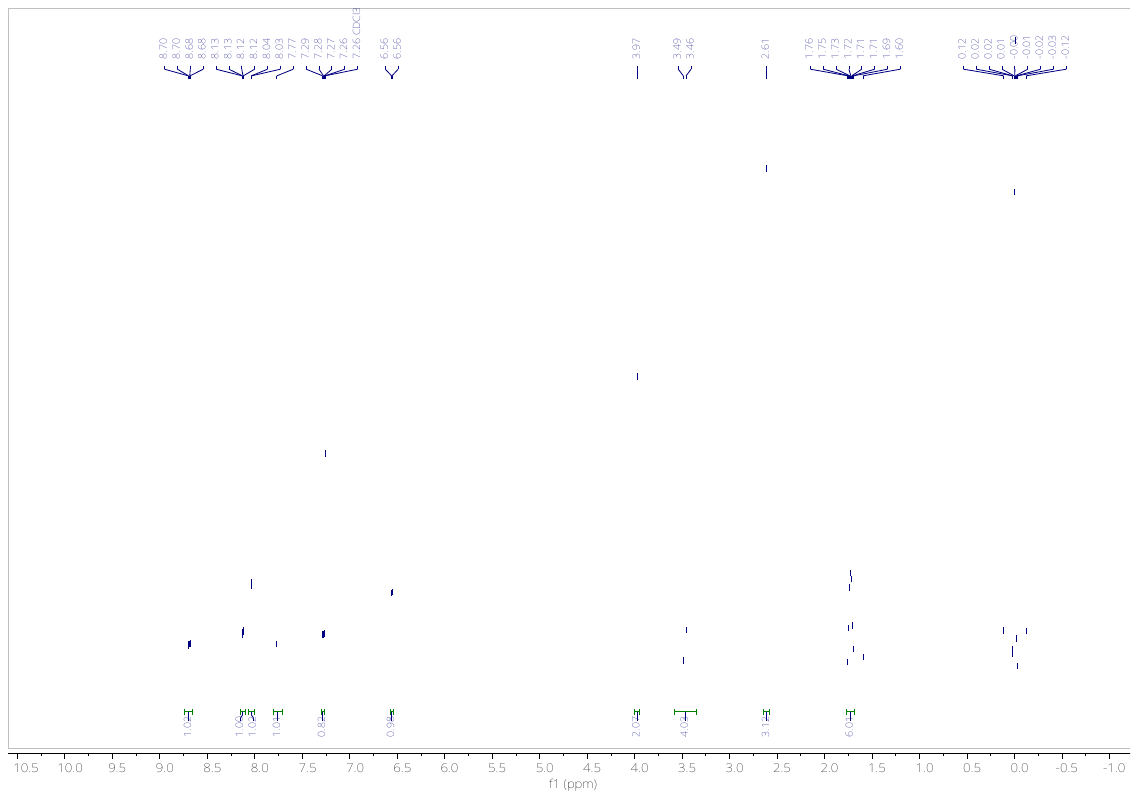

**Fig. 7.** ^1^H NMR of **EN11**.

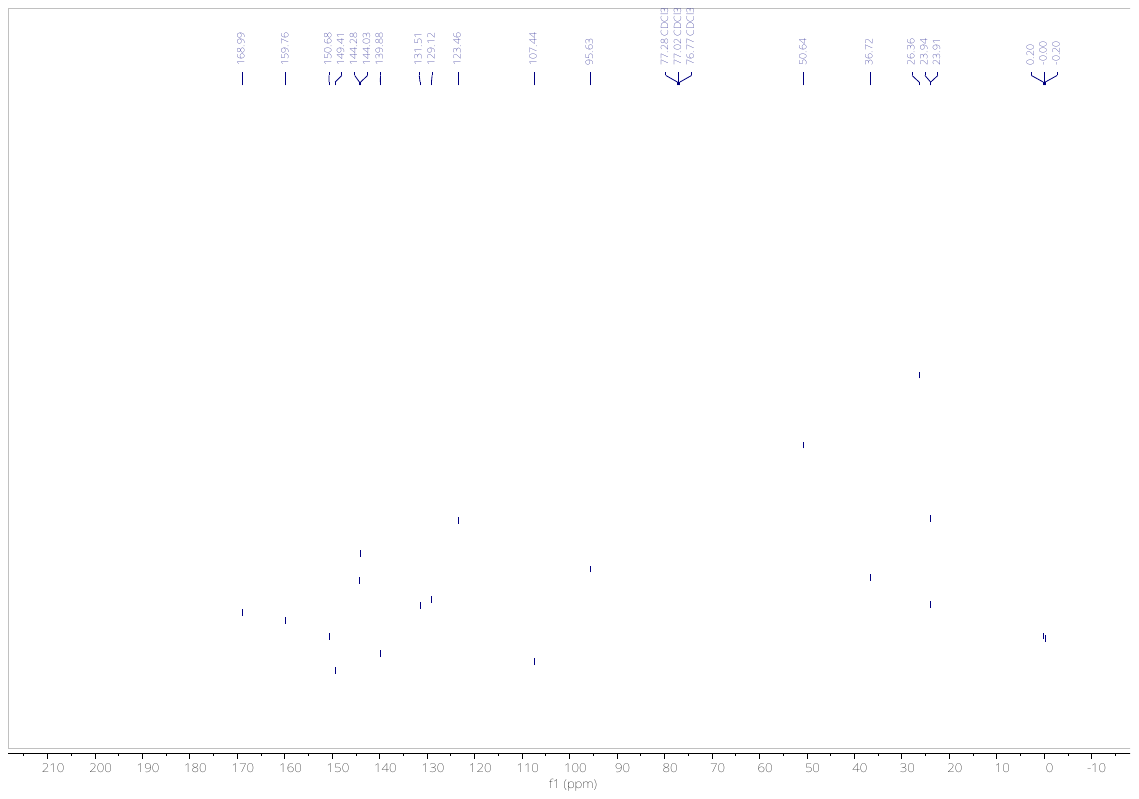

**Fig. 8.** ^13^C NMR of **EN11**.

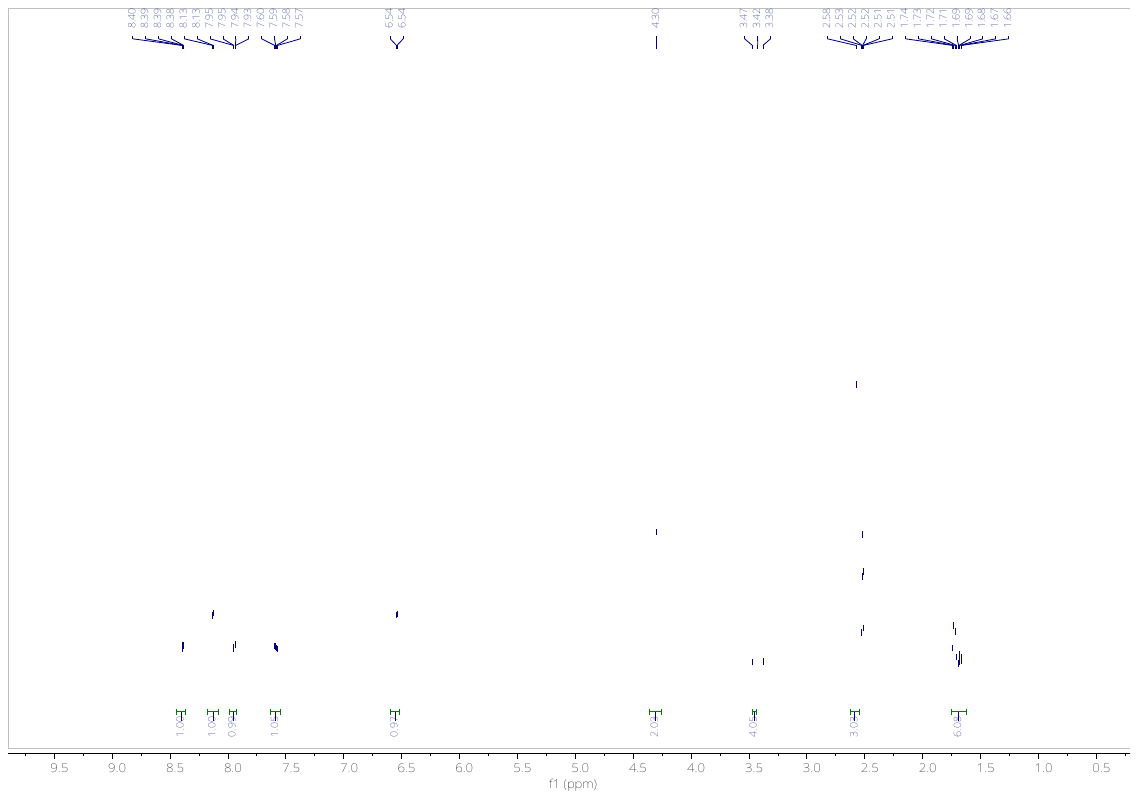

**Fig. 9.** ^1^H NMR of **EN12**.

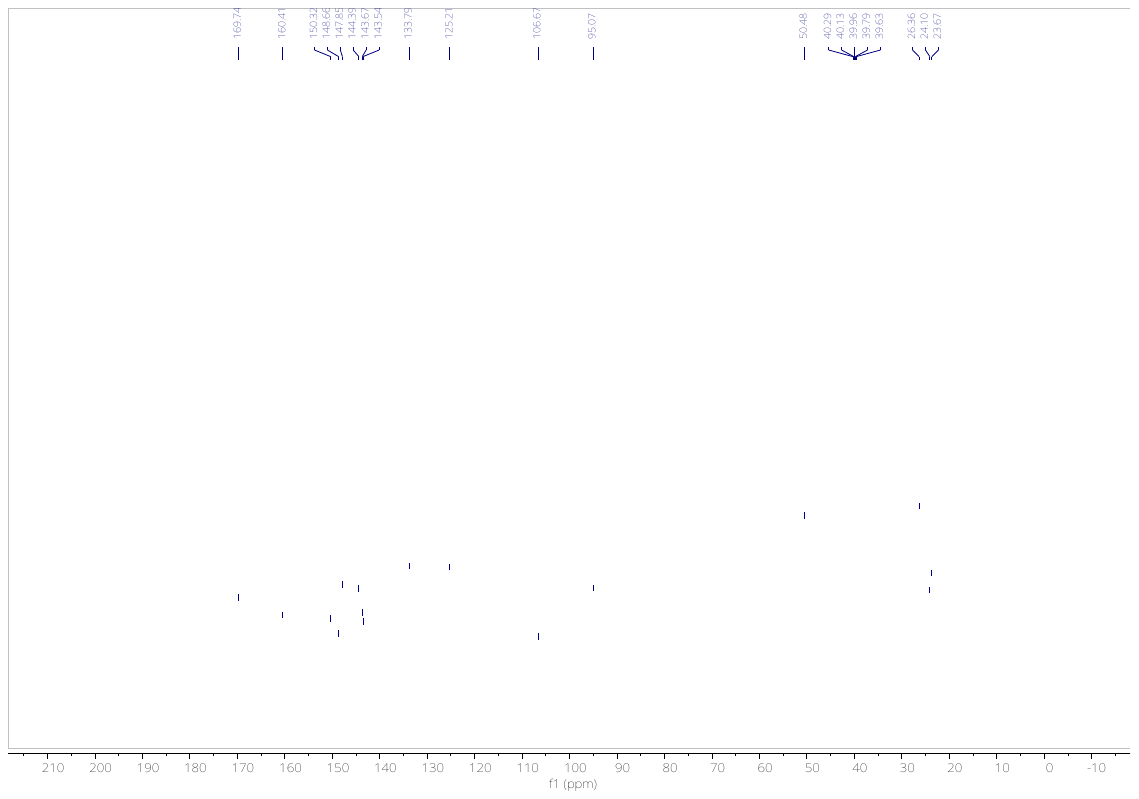

**Fig. 10.** ^13^C NMR of **EN12**.

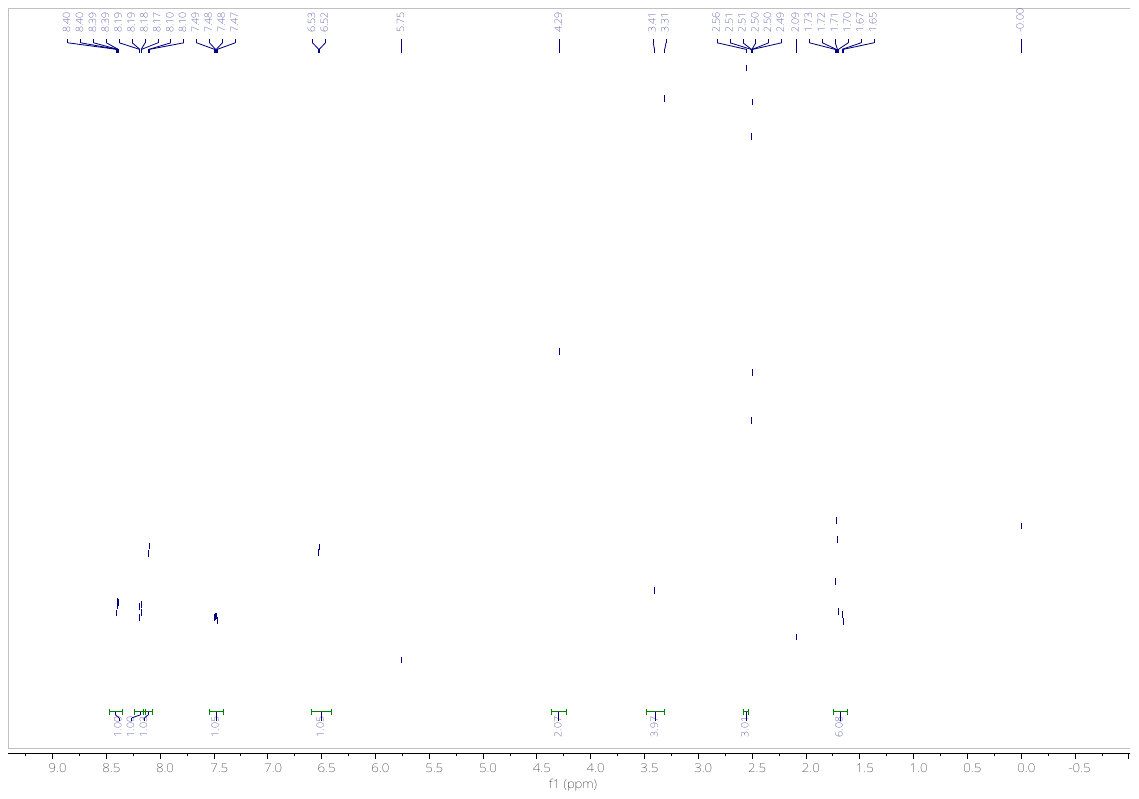

**Fig. 11.** ^1^H NMR of **EN13**.

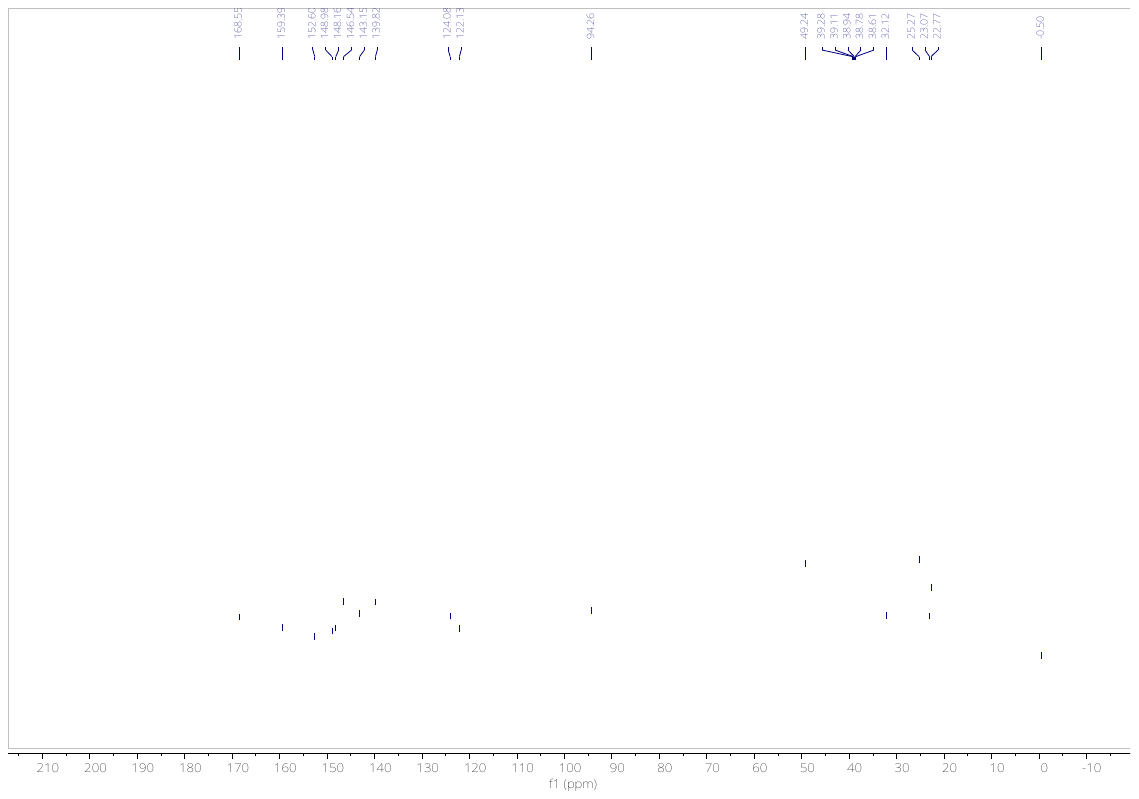

**Fig. 12.** ^13^C NMR of **EN13**.

**HRMS Spectra:**

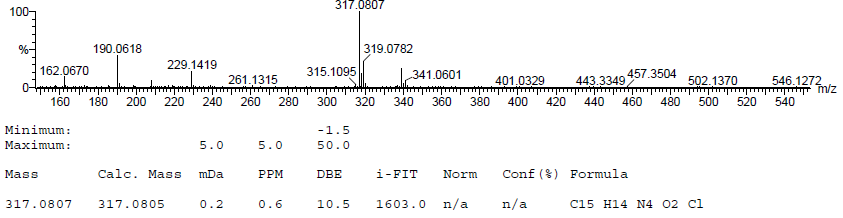

**Fig. 13.** HRMS of **EN5.**

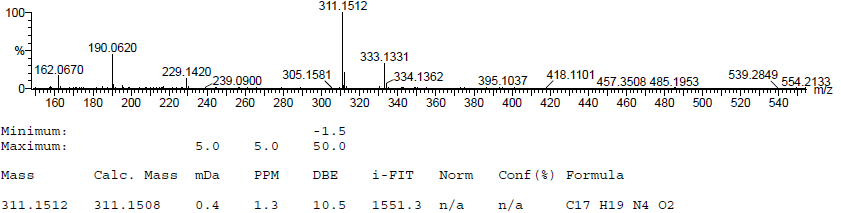

**Fig. 14.** HRMS of **EN6.**

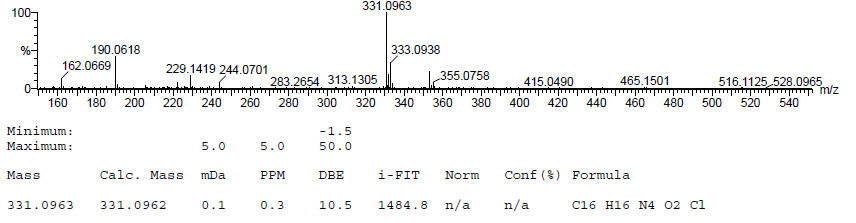

**Fig. 15.** HRMS of **EN7.**

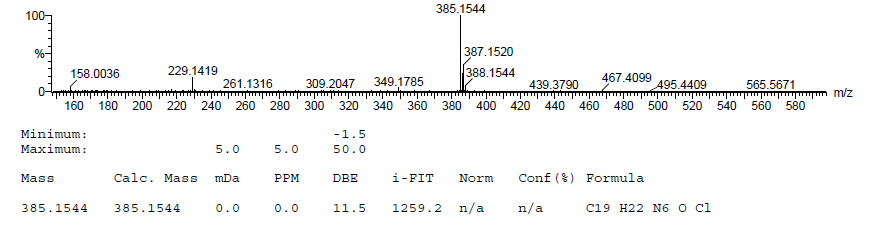

**Fig. 16.** HRMS of **ENP11.**

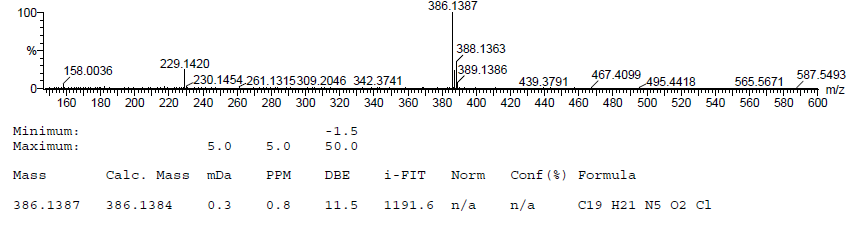

**Fig. 17.** HRMS of **ENP12.**

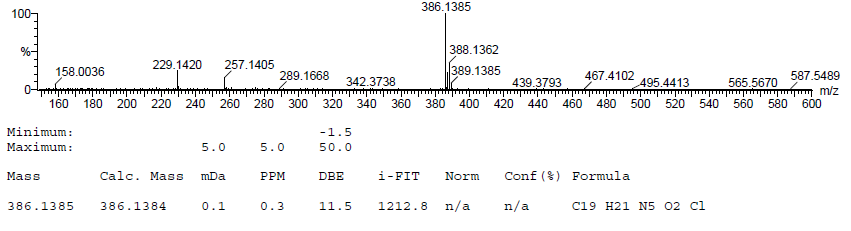

**Fig. 18.** HRMS of **ENP13.**
